# Metagenomics reveals niche partitioning within the phototrophic zone of a microbial mat

**DOI:** 10.1101/151704

**Authors:** Jackson Z Lee, R Craig Everroad, Ulas Karaoz, Angela M Detweiler, Jennifer Pett-Ridge, Peter K Weber, Leslie Prufert-Bebout, Brad M Bebout

## Abstract

Hypersaline photosynthetic microbial mats are stratified microbial communities known for their taxonomic and metabolic diversity and strong light-driven day-night environmental gradients. In this study of the upper photosynthetic zone of hypersaline microbial mats of Elkhorn Slough, California (USA), we show how reference-based and reference-free methods can be used to meaningfully assess microbial ecology and genetic partitioning in these complex microbial systems. Mapping of metagenome reads to the dominant *Cyanobacteria* observed in the system, *Coleofasciculus (Microcoleus) chthonoplastes*, was used to examine strain variants within these metagenomes. Highly conserved gene subsystems indicate a core genome for the species, and a number of variant genes and subsystems suggest strain level differentiation, especially for carbohydrate utilization. Metagenome sequence coverage binning was used to assess ecosystem partitioning of remaining microbes. Functional gene annotation of these bins (primarily of *Proteobacteria, Bacteroidetes,* and *Cyanobacteria*) recapitulated the known biogeochemical functions in microbial mats using a genetic basis, and also revealed evidence of novel functional diversity within the *Gemmatimonadetes* and *Gammaproteobacteria*. Combined, these two approaches show how genetic partitioning can inform biogeochemical partitioning of the metabolic diversity within microbial ecosystems.

## Introduction

Hypersaline microbial mats are diverse laminated assemblages of microorganisms thought to represent one of the earliest ecosystems on Earth, and are typically dominated by oxygenic phototrophic cyanobacteria (Awramik, 1984; Oren 2010). Compact and highly structured, these mats contain microbial communities that possess great diversity at both the metabolic and phylogenetic level (Canfield & Des Marais 1993; Ley et al. 2006). Microbial mats have been described as complete ecosystems in miniature, with relatively closed cycling of photosynthetically fixed carbon from the upper layers distributed to heterotrophic organisms in the lower layers for re-mineralization and subsequent reincorporation. Photosynthetic activity of oxygenic phototrophs during the daytime is followed by a rapid transition into anoxic conditions following sunset. Features of an active nitrogen cycle include high rates of nitrogen fixation supported by daytime photosynthetic activity or sulfide redox reactions (Bebout et al. 1993).

Extensive biogeochemical, microbiological, and targeted molecular ecological studies have been completed on the hypersaline microbial mats of Elkhorn Slough (CA) and have identified rates of biogeochemical processes and the identities of some organisms involved. As an example, previous studies have shown that within the upper 2 mm layers of these mats, net hydrogen production is a consequence of constitutive fermentation of photosynthate to acetate by *Cyanobacteria,* followed by consumption of fermentation byproducts by *Desulfobacterales* and *Chloroflexi* (Burow et al. 2012, 2013, 2014; Lee et al. 2014). Major nitrogen fixers have also been identified, and include a novel group of cyanobacteria (ESFC-1), that have been isolated and whole genome sequenced (Woebken et al. 2012; Stuart et al. 2015; Everroad et al. 2016). In many cases, identity and metabolic role of microbes have not been linked to specific biogeochemical transformations, largely due to the high diversity and novelty of the microorganisms present in these mats.

In this study, high throughput shotgun metagenomic sequencing was combined with several genome-centric bioinformatics approaches (co-assembly, coverage binning, reference-mapping) to understand functional and microbial diversity in the phototrophic zone of microbial mats of Elkhorn Slough, CA. Underpinning this work are previous binning studies have sought to analyse metagenomic results at the organism level rather than at the microbiome level, either to identify novel genomic diversity (Dick et al. 2009; Wrighton et al. 2012; Hanke et al. 2014; Sekiguchi et al. 2015), novel metabolism (Podell et al. 2013), novel genetics (Hess et al. 2011; Nielsen et al. 2014), or ecological succession (Morowitz et al. 2011; Brown et al. 2013; Sharon et al. 2013). Using reference-free binning of metagenomic scaffolds we show that partitioning of metagenomes derived from microbial mats resulted in recovery of numerous bins representing species, or groups of closely related organisms, in the phototrophic mat layer. Using reference-based variant analysis with a sequenced genome of the dominant *Cyanobacteria Coleofasciculus chthonoplastes*, we identified genes representing metabolic pathways conserved within this species. Analysis of the functional genetic diversity of these dominant bins recapitulated the known partitioning of ecological functions in these mats and also identified novel organisms that have yet to be isolated. **This combined reference and reference-free based approach shows the value of binning metagenomes and provides a robust systems biology analysis of novel microbial ecosystems from a genetic perspective.**

## Methods

### Site sampling and incubation description

Samples were collected from the Elkhorn Slough estuary at 36°48’46.61” N and 121°47’4.89” W. The site consists of up to 1 cm thick mats dominated by *Coleofasciculus* sp. (formerly *Microcoleus* sp.) and *Lyngbya* sp., that vary with seasonal water flows and nutrient inputs. The conditions and the mats found at this site have been documented in previous reports (Burow et al. 2012, 2014; Woebken et al. 2012). A single contiguous mat piece approximately 60-80 cm in diameter was harvested in a total of thirty-two 10 cm diameter acrylic cores tubes on Nov. 8, 2011 at 6 AM. Cores were sealed with rubber stoppers on the bottom, covered with clear plastic wrap and transported to NASA Ames Research Center (Moffett Field, CA) for incubation with Elkhorn Slough water in aquaria under natural light (Figure S1). The cores were split into two treatments: controls, and a set treated with 30 mM molybdate to inhibit sulfate reduction. Cores were incubated in a temperature controlled water bath, monitored, and sacrificed in 4-hour intervals over 24 hours. Two control and two molybdate-treated samples, collected at 0130 and 1330, were selected for metagenomic sequencing. Because of the short timeframe, no differences in community composition were expected amongst these samples.

### Nucleic acid extraction

Nucleic acid extraction was conducted using a total RNA/DNA approach consisting of a phenol-chloroform method described by Woebken et al. (2012) in combination with the RNeasy Mini Elute Cleanup Kit and QIamp DNA Mini Kit (Qiagen, Hilden, Germany). A rotor-stator homogenizer (Tissue-master, Omni International, Kennesaw, GA, USA) was first washed successively with 70% EtOH, RNase-away (Sigma, St. Louis, MO, USA), and RNase-free H_2_O. The top 2 mm of the microbial mat was excised using a sterile razor blade and homogenized for 30 seconds on the lowest setting in 0.5 ml RLT buffer mix (10 ml RLT buffer (RNEasy Plus Mini kit, Qiagen) and 100 ul β-mercaptoethanol) in a 2 ml bead beating tube (0.5 mm zirconium beads). A FastPrep bead beater (MP Biomedicals, Santa Ana, CA, USA) was used for 40 seconds at setting “6.0”. Samples were spun for 1 minute at 8,000 x *g* (rcf) and the supernatant transferred to new 2 ml tubes. DNA was isolated by adding an equal volume of phenol-chloroform (basic) and vortexing for 10 seconds, incubating for 5 minutes at room temperature, and spinning for 5 minutes at 8,000 x *g* (rcf). The aqueous phase was transferred to a new tube on ice. An equal volume of 100% ethanol was added to eluate and vortexed for 10 seconds. 700 ul of supernatant/ethanol mix was added to QIAmp spin column (QIAamp DNA mini prep kit, Qiagen) and processed according to manufacturer’s instructions. RNA was also extracted from each sample using RNA Mini Prep kits (Qiagen) and reserved for future gene expression analysis. Whole genome shotgun metagenomic sequencing was completed at the Joint Genome Institute (JGI) on an Illumina HiSeq 2000 platform (Lee et al., in submission).

### Read preprocessing, assembly, and annotation

150 bp paired end reads were quality trimmed using Trimmomatic (Bolger et al. 2014) with the following parameters: headcrop 11, trailing 20, sliding window 4:20, minimum length 75 bp. Quality filtered reads were assembly with Ray-Meta (Boisvert et al. 2012) on the NERSC Edison supercomputing cluster. Sequences from metagenomes of microbial mat samples were pooled together and co-assembled with Ray-Meta (Boisvert et al. 2012) three times using different assembly word sizes (k=29, 45, 63). Assembly word sizes were assessed using KmerGenie (Figure S2A) (Chikhi & Medvedev 2014). Each individual metagenome sample was then mapped using Bowtie2 (Langmead & Salzberg 2012) back to each assembled scaffolds to determine sample specific coverage. Prodigal (Hyatt et al. 2010) was then used to predict open reading frames (ORFs), and an HMM model (Eddy 2011) was used to find the essential single copy genes (Albertsen et al. 2013). All ORFs were submitted to MG-RAST (Meyer et al. 2008) for annotation using a BLAT 90% clustering protocol. The mapping process was repeated for open reading frames (ORFs) detected by Prodigal (Hyatt et al. 2010). MG-RAST custom md5 matches were quality filtered, de-replicated, and parsed to a tab-delimited ORF database of ORF name, protein annotation, ontology annotation, taxonomy, and coverage.

### Metagenomic binning

Previous work examining binning has shown several successes using k-mer nucleotide frequency, %GC, read coverage, taxonomy, or a combination of these strategies as biosignatures of genomes within metagenomes. The pipeline used in this study combines supervised learning (Dick et al. 2009), dimensionality reduction (Woyke et al. 2006), and the coverage binning (Albertsen et al. 2013) using R and CRAN analysis packages.

As was noted by Dick et al. (2009), preliminary results suggested that larger scaffolds harbored strong phylogenetic signal (Figure S2B), so these scaffolds were used to recruit clusters representing bins from the metagenomes. Log normalization and principle component analysis (PCA) dimensionality reduction using scaled values of %GC and differential sample coverage aided to resolve binning ‘spears’ seen in metagenomic data. DBScan, noted for being sensitive to cluster density and used on noisy datasets (Ester et al. 1996), was selected to cluster large (>5 kbp) scaffolds in PCA space. The remaining scaffolds >1.5 kbp were recruited using a SVM machine-learning algorithm (Chang & Lin 2011) trained on the larger scaffolds (Supplemental Figure S3B) and was tuned by maximizing single copy essential gene membership and minimizing gene copy duplication. This was repeated for the 3 assemblies (performed at different word sizes (k=29, 45, 63)); the best corresponding bin (maximum single copy genes, minimum duplication) from each assembly was extracted and pooled with both background and unbinned contigs from the k=29 assembly. Binning procedure and quality analysis were based on analysis of ~100 essential single copy genes (Albertsen et al. 2013; Dupont et al. 2012). These data when charted together produce informationally-dense PCA graphs overlaying the clustering of contigs with contig information (taxonomy, contig size, etc.) which we refer to as “galaxy” plots.

### Annotation search and collation

Since a number of different ontology systems and gene annotation databases were used to annotate genes, querying single annotation sets (e.g. KEGG) produced partial results. To maximize search coverage, a customized Python regular expression search algorithm was developed. This algorithm emphasized matching KEGG and EC ontologies (e.g. K02586 and 1.18.6.1, with subunit or chain designation), but can also search for multiple text patterns in protein annotations by keyword and subunit (e.g. nitrogenase alpha chain), and protein abbreviation (e.g. nifD). The algorithm also allows for nested searches (e.g. first cytochrome c oxidase, then cbb3 subtype), as well as exclusion terms (e.g. ‘precursor’ proteins). Results were cross-referenced with bin annotation and written to tab-delimited files for heatmap generation in Excel. A list of genes associated with biogeochemical cycling (sulfur metabolism, nitrogen metabolism, phototrophy, autotrophy, and hetrotrophy) and with starch utilization were determined and used to query annotation records.

### Read mapping and variant analysis

Recent studies have examined the possibility of using variant callers typically seen in human genomic variant analysis for the detection of strain variation across genomes (He et al. 2010) and for differing populations of *Bacteria* in the human gut microbiome (Schloissnig et al. 2013). We used these insights and approach to call the coverage and density of variants in genes and in subsystems that differed from the mapping reference. The complete metagenomic dataset from the selected four mat samples were pooled and mapped to the *C. chthonoplastes* PCC 7420 (Rippka et al, 1979, Garcia-Pichel et al. 1996) genome (GCA_000155555.1 ASM15555v1, JCVI) using Bowtie2 with default settings. Single nucleotide polymorphisms (SNPs) were called with FreeBayes using haploid continuous pooled variant calling settings (Garrison & Marth 2012). Variants were filtered for poorly called variants and selected for SNPs using Samtools and BCFtools (Li et al. 2009; Danecek et al. 2011). Bedtools (Quinlan & Hall 2010) was used to count the number of variants per gene in the PCC7420 genome. Variants were summarized by gene and SEED subsystems using PATRIC annotations (Wattam et al. 2013) cross-referenced with NCBI annotations (with a cutoff >70% annotation overlap using Bedtools). SNPs were then filtered for coverage between 50-200 reads and the ratio of SNPs per all bases in a gene (SNP density) was calculated. These gene SNP density values were matched to subsystem ontology. Each gene was binned by SNP density into 4 groups, (0-1%, 1-2%, 2-3%, 3%+) variants / gene length. A score was developed based on the aggregation of gene variant density in subsystems to estimate the level of genetic variation in each subsystem as compared to genome-wide variation levels (Equation 1)
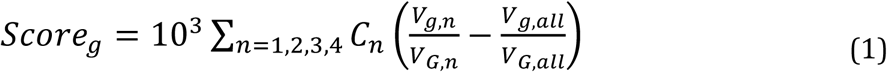

where: *g* is a subsystem gene set from all gene sets *G*, *n* is the variant density bin described above, and *V_g,n_* is the number of genes in set *g* also in bin *n*, and *C_n_* is a weighting coefficient for each bin (here, *C_n_*= 1, unweighted). A variation ratio of *V_g,n_* to number of genes in *n* for *G* (*V_G,n_*) was calculated and offset by the variation ratio of *all* gene bins in subsystem *g* and then summed for each bin. The result was scaled by an arbitrary factor of 1,000 for convenience to generate a final score for each gene set *g*. Positive scores indicated more genes with variation in a subsystem than the genome-wide average, negative scores indicated fewer genes with variation than the genome-wide average.

### Data and repository archival

Data and code from this study were archived to several locations. JGI sequences were archived on the JGI IMG server under Project ID 1081546-1081548,1000633. Ray-meta assemblies and gene annotation tables were archived at 10.5281/zenodo.584152. All codes used on this project are available at GitHub at (http://github.com/leejz/). Three-dimensional interactive ShinyRGL visualizations of galaxy plots are available at https://leejz.shinyapps.io/plot3d2 (training dataset) and at https://leejz.shinyapps.io/plot3d3 (full dataset).

## Results

### Metagenomic binning of the photosynthetic zone of microbial mats

Initial analysis was performed at the JGI on all four metagenomes. Coverage and diversity estimation indicated essentially equivalent metagenome composition at a phylum level (Lee, et al., in submission) and suggested that metagenomes could be combined for differential coverage analysis. “Galaxy” charts of PCA components overlaid with taxonomic information from essential single copy genes found on contigs (Figure 1A) showed that bins could be identified from longer contig fragments containing these genes. When longer contigs were clustered, bins were clearly delineated as density-dependent clusters by the first 3 principal component axes (Supplemental Shiny visualization plots). This produced more than 70 bins (Figure 1B) evaluated for completeness (fraction of essential single copy genes detected), duplication (fraction of duplicate essential single copy genes detected) of which the top 20 bins (equivalent to those with a completeness > 80%) were selected for downstream analysis. Table 1 shows the top 20 ordered by mean coverage and Supplemental Dataset 1 shows the remainder. Two approaches to determine taxonomic affiliation were undertaken. In a more conservative approach, we selected the most common phylum annotated by Hidden Markov Model (HMM) profiles of essential single copy genes. Where taxonomy could be determined for these single copy genes, taxonomy at phylum level was concordant for each bin, with one exception (Table 1, bin 6). Our second method for taxonomy determination was to examine the most common genome annotation matched by MG-RAST for all Open Reading Frames (ORFs) in a bin. This produced a nearest genome result and the fraction of genes that matched to this nearest genome (Table 1). Of the 20 bins selected for downstream analysis, using annotations of single copy essengial genes, 3 bins of *Cyanobacteria* (bins 1,2,9), 5 bins of *Gammaproteobacteria* (bins 3,4,7,8,20), 1 bin of *Alphaproteobacteria* (bin 10), 1 bin of *Deltaproteobacteria* (bin 16), 1 bin of *Firmicutes* (bin 6), and 9 bins of *Bacteroidetes* (bins 5,11,12,13,14,15,17,18,19) were annotated with near consensus at the phylum level (class level for *Proteobacteria*). Consensus species taxonomic annotations from ORF annotation of the most abundant two bins (bins 1, 2) suggested that the dominant *Cyanobacteria* known from these mats (*C. chthonoplastes*, and *Lyngbya* sp.) were captured. The L50 assembly metric for bin 1 (*C. chthonoplastes*) was much lower than expected (despite the fact that this taxa was the most abundant and therefore had the deepest read sampling). The taxonomy of the third most abundant bin (bin 3) suggested that this was a purple sulfur bacteria (*Thiorhodovibrio* sp.); the remaining *Gammaproteobacteria* (bins 4,7,8,20) were poorly matched to reference genomes. The lone *Firmicutes* bin (bin 6) was unique in that there was no strong consensus of phylum identity among annotated essential-copy genes, and it had ORF annotations that belonged to the only sequenced member of the phylum *Gemmatimonadetes*. The single bin of *Alphaproteobacteria* (bin 10) in this subset was annotated as a possible relative of *Rhodospirillum rubrum* (54% of ORFs). Many bins in this subset annotated to various *Bacteroidetes*. Several of these bins (bins 11,13,15,19) had a closest genome match to a beach sediment chemoheterotroph, *Marivirga tractuosa* DSM 4126 (Pagani et al. 2011), but only one of these (bin 19) shared a large amount of similarity (96% of ORFs). One bin (bin 14) matched to *Psychroflexus torquis* ATCC 700755 (Bowman et al. 1998) (80% of ORFs) derived from Antarctic ice.

**Figure 1.**
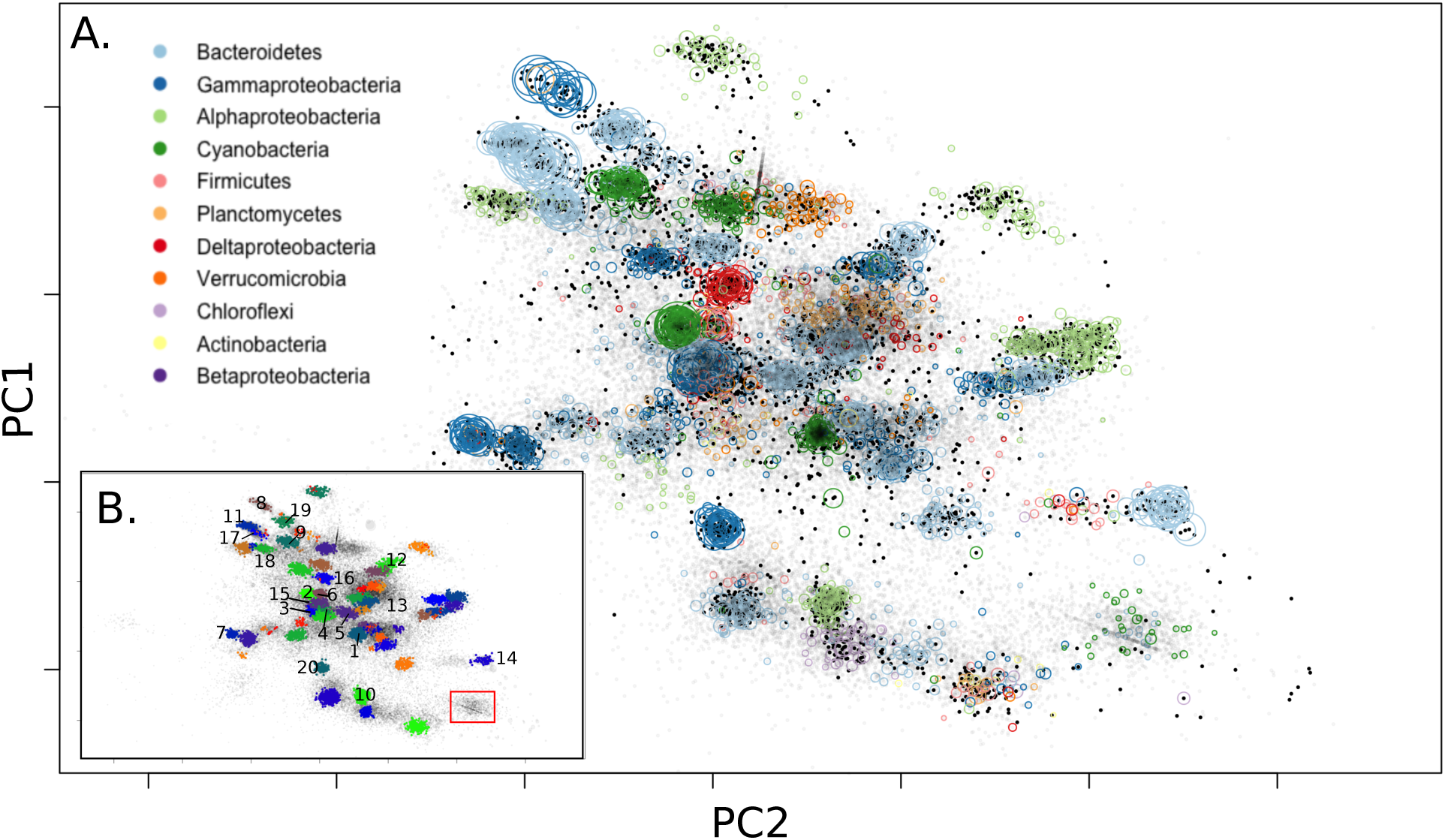
PCA galaxy chart with MG-RAST annotated abundant phyla labeled (A). Dark dots represent the >5kbp training dataset, and light dots represent all scaffolds >1.5kbp. Colored circles represent phylum of contigs based on single copy essential gene classification. Size of phylum circles is proportional to contig size. The top two axes are shown here, but the third largest component was also used to differentiate bins. Final detected bins are shown in (B) with complete bins from Table 1 enumerated and Cyanobacterium ESFC-1 labeled (red box).

**Table 1.**
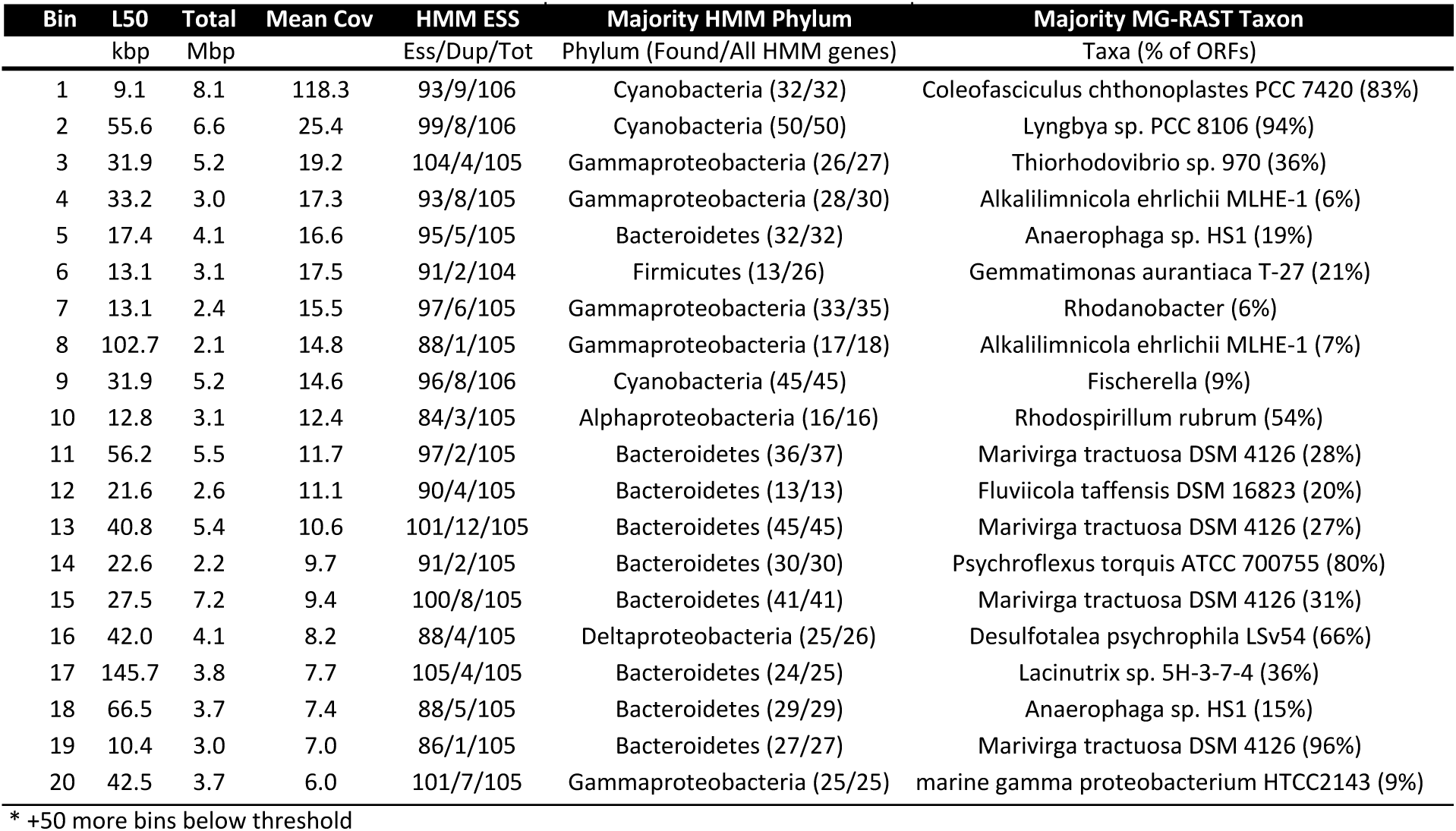
Summary statistics for each bin: scaffolds (L50: L50 contig size, Total: total Mbp binned, Mean Cov: mean coverage of all scaffolds), HMM essential single copy gene completeness (ESS: Essential single copy genes, Dup: duplicated ESS, Tot: All ESS in phylum), majority HMM phylum: majority of identified taxonomy in ESS genes, and majority MG-RAST Taxon: most common genome identified in ORFs.

A full listing of all bins identified in this study is included in Supplemental Dataset 1 with a cross-referenced table (Figure S3A) for each assembly. We note that some of the minor bins, still largely unexamined, contained annotations for possible *Chloroflexi*, *Planctomycetes*, and *Verrucomicrobia* and may prove useful to future studies of the diversity of these organisms.

### Bin pathway annotations comparing *C. chthonoplastes* to other *Cyanobacteria*

Binning and annotation analyses showed bins 1 and 2 closely matched to *C. chthonoplastes* PCC 7420 and *Lyngbya* sp. PCC 8106 respectively. A third bin (bin 9) loosely matched a *Fischerella* genome (9% of ORFs). To understand the metabolic differences between *Cyanobacteria* within mats, annotations from these 3 bins were combined with annotations from the draft genome of ESFC-1, a nitrogen-fixing cyanobacterium previously isolated from the same ecosystem and sequenced (Woebken et al. 2012; Everroad et al. 2016). In the current study ESFC-1 did not pass binning thresholds, although it was detected in metagenomes (Figure 1B). To draw comparisons between these, the major KEGG pathways were visualized. Notable differences were observed in the carbohydrate utilization pathways (Figure S4). When annotations of ESFC-1 were included as a bin with the 3 other *Cyanobacteria* and explored in depth by searching for pathway gene terms in annotations, the *C. chthonoplastes* bin had a unique set of glycoside hydrolases involved in many aspects of both carbohydrate production and breakdown (GH3, GH5, GH9, GH38, GH57, Figure 2A). All bins contained annotations for several types of cellulases, cellulose synthesis genes, and starch (glycogen) storage and had some involvement of maltose and sucrose synthesis and cycling (Figure 2B). A full table of all search terms used and all extracted search results are available in Supplemental Dataset 1.

**Figure 2.**
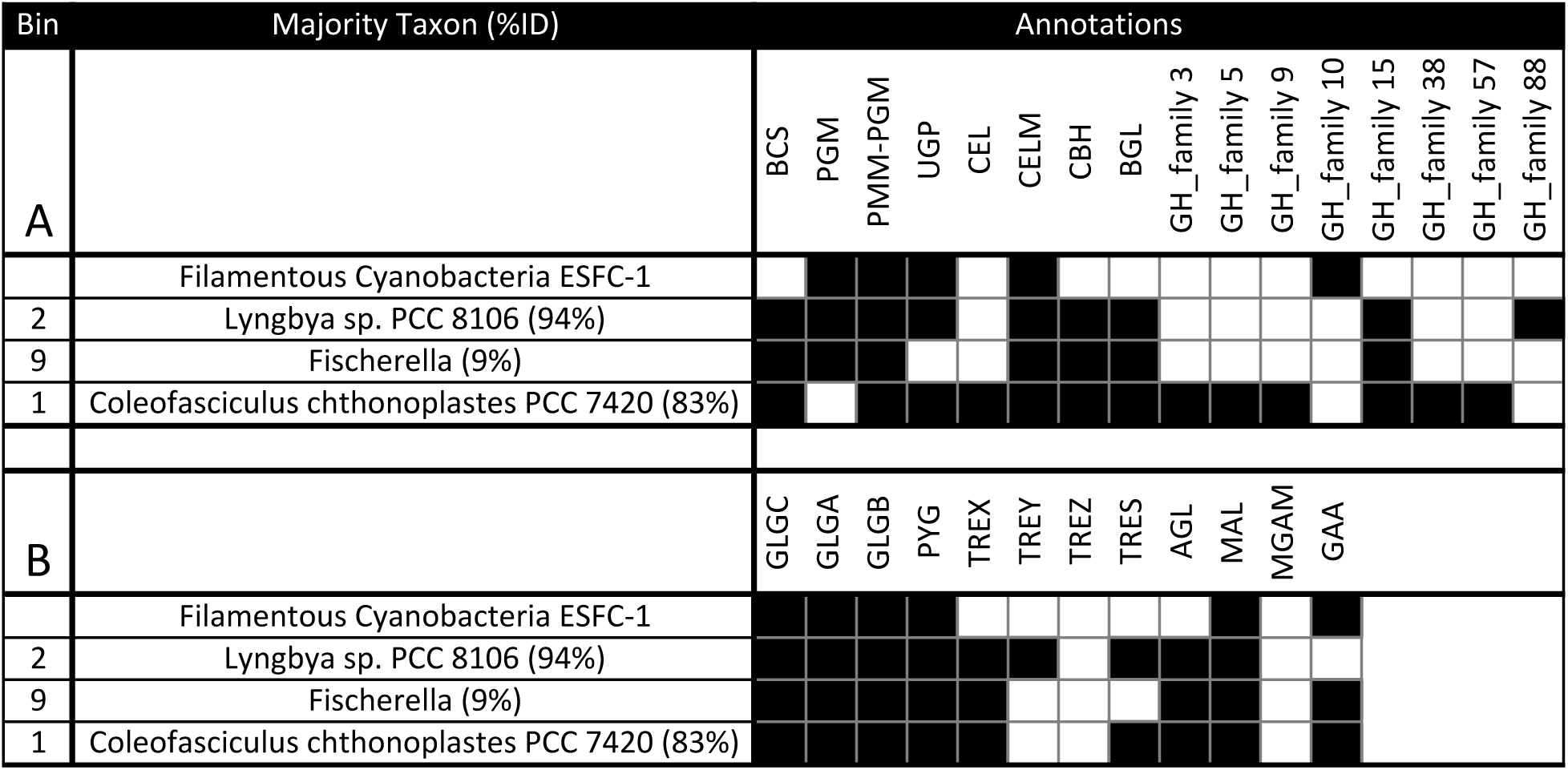
Carbohydrate regulation from *Cyanobacteria* bins and the genome of ESFC-1 indicate unique polysaccharide capability of *C. chthonoplastes* among mat organisms when examining cellulose production genes (A), starch production genes (B). Abbreviations: BCS: cellulose synthase, PGM: phosphoglucomutase, PMM-PGM: phosphomannomutase/phosphoglucomutase, UGP: UTP--glucose-1-phosphate uridylyltransferase, CEL: Cellulase / Endoglucanase, CELM: cellulase M, CBH: cellulose 1,4-beta-cellobiosidase, BGL: beta-glucosidase, GH: glycosyl hydrolase, GLGC: glucose-1-phosphate adenylyltransferase, GLGA: glycogen synthase, GLGB: 1,4-alpha-glucan branching enzyme, PYG: glycogen phosphorylase, TREX: glycogen operon protein, TREY: maltooligosyltrehalose synthase, TREZ: maltooligosyltrehalose trehalohydrolase, TRES: trehalose synthase, AGL: glycogen debranching enzyme, MAL: 4-alpha-glucanotransferase, MGAM: maltase-glucoamylase, GAA: alpha-glucosidase

### Functional genetic diversity of biogeochemical cycling identified using metagenomic bin annotations

A catalog was compiled of the pathways involved with biogeochemical cycling (C, N, S) and of the organisms involved with those pathways. Briefly, annotations of indicator genes selected from these pathways were used to identify the partitioning of biogeochemical roles across bins as well as the remaining ecosystem (minor bins, background recruitment bin, and remaining unbinned genes (Figure 3). A complete list of bins, genes used in this study, gene abbreviations used in this study, and gene selection criteria can be found in Supplemental Dataset 1. Clear delineations between bins involved in sulfur cycling, nitrogen cycling, and carbon cycling could be observed. For example, both sulfur oxidation and sulfate reduction could be resolved. Capacity for phototrophic sulfur oxidation, represented by DSR, APR, and SOX genes for sulfur oxidation and PUF, CHL, BCH for bacterial phototrophy, was present in a bin (bin 3) representing the dominant clade of *Thiorhodovibrio* sp. A *Deltaproteobacteria* (bin 16) was annotated as sulfate-reducing, represented by DSR, APR, and methyl viologen-reducing hydrogenase (MVH). This bin also contained a suite of annotations for oxygen tolerance such as reactive oxygen scavenging indicated by rubrerythrin, thioredoxin, catalase-peroxidase, and alkyl hydroperoxide reductase, as well as direct oxygen scavenging indicated by rubredoxin, molybdopterin oxidoreductase, NADH-quinone oxidoreductase (Supplemental Annotations). A bin associated with purple non-sulfur bacteria (PNS) (bin 10) was also identified as having SOX genes.

**Figure 3.**
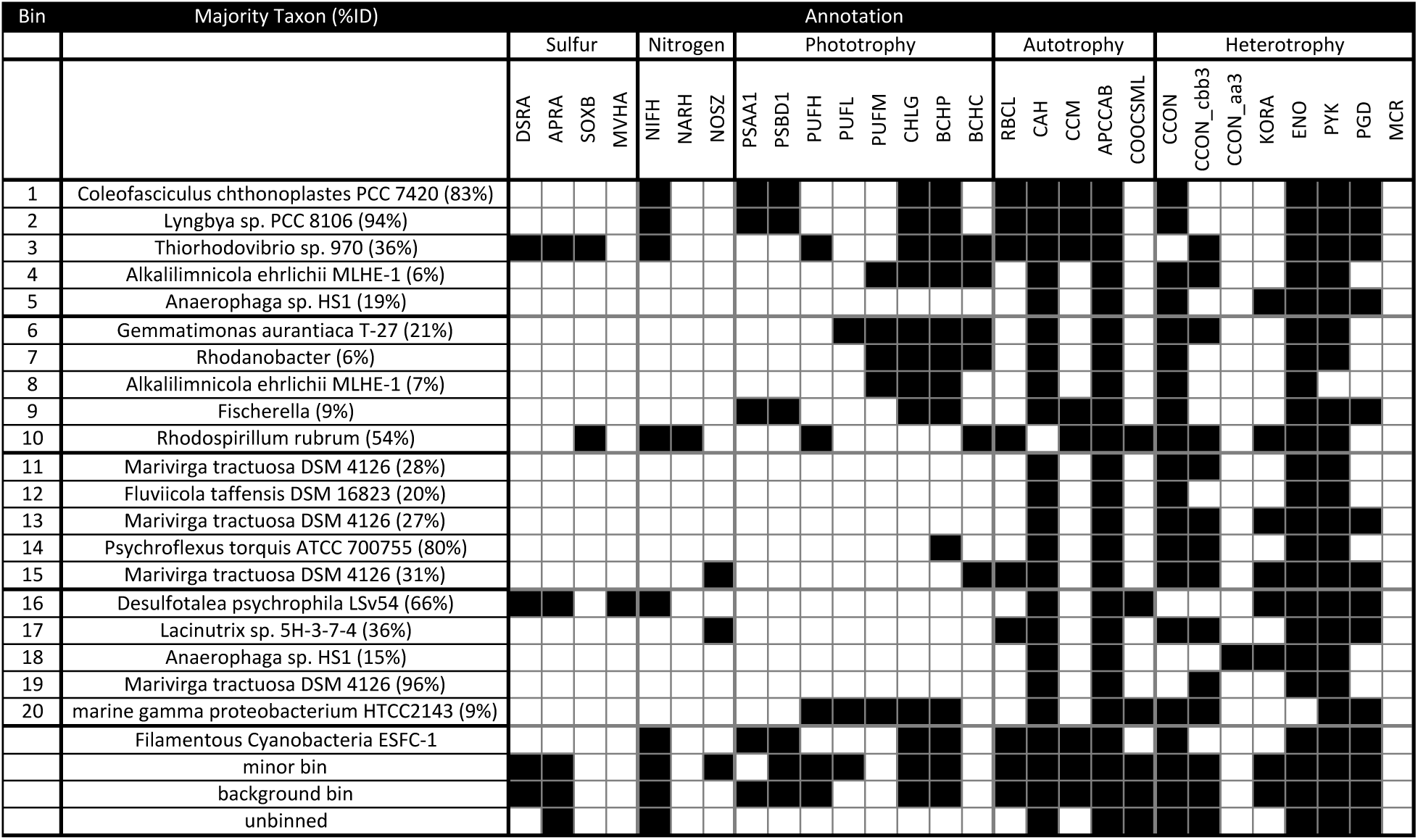
Summary table of annotated chlorophyll types and putative metabolism, one row for each major bin and one representative gene per column. Also included are annotations from Cyanobacterium ESFC-1, minor bins, background bin, or unbinned scaffolds. Each label includes a three-letter abbreviation, and subunits examined (e.g. DSRA: Dissimilatory sulfate reductase A). (Abbreviations: DSR: dissimilatory sulfite reductase, APR: adenylylsulfate reductase, SOX: sulfite oxidase, MVH: methyl viologen-reducing hydrogenase, NIF: nitrogenase, NAR: nitrate reductase, NOS: nitrous-oxide reductase, PSA: photosystem I P700 chlorophyll a apoprotein A1, PSB: photosystem II protein D1;photosystem II protein D2, PUF: photosynthetic reaction center, PSC: photosystem P840 reaction center, CHL: chlorophyll synthase;bacteriochlorophyll a synthase, BCH: bacteriochlorophyll c synthase, RBC: ribulose bisphosphate carboxylase, CAH: carbonic anhydrase, CCM: carboxysome microcompartment protein, APCC: acetyl-CoA carboxylase and/or propionyl-CoA carboxylase, COO: carbon monoxide dehydrogenase, CCON: cytochrome c oxidase, KOR: 2-oxoglutarate synthase, ENS: enolase, phosphopyruvate hydratase, PYK: pyruvate kinase, PGD: 6-phosphoglucanate dehydrogenase, MCR: methylcoenzyme M reductase)

Within the nitrogen-utilizing bins, annotations for nitrogen fixation (NIF) were detected in 5 bins (bins 1,2,3,10,16), and in minor, unbinned, and background bins, suggesting that this essential trait was widespread. However, nitrate reduction was observed in only one bin (PNS bacteria, bin 10) and not in minor or background bins. Likewise denitrification was only observed in two bins (bins 15, 17).

Within carbon-cycling annotations, several types of phototrophy could be observed. Photosystem I/II photoautotrophy represented by PSA, PSB, CHL, RubisCO (RBC), carbonic anhydrase (CAH), carboxysome microcompartment protein (CCM), and acetyl/propionyl-CoA carboxylase (APCC) was found in several *Cyanobacteria* bins (bins 1,2,9). The one PNS bin (bin 10) had annotations for bacteriochlorophyll-based photoautotrophy, represented by PUF, BCH, RBC, CCM, and APCC, and carbon monoxide dehydrogenase (COO). A number of PNS bins were also noted in the minor bin dataset, in this case annotating as relatives of *Erythrobacter* sp. (Supplemental Dataset 1). In addition to these known phototrophs, we also noted a number of other bins contained phototrophic genes. Four bins (bins 4,7,8,20) annotated as novel *Gammaproteobacteria* contained bacterial phototrophic genes, represented by PUF, CHL, and BCH. Additionally, bin 6, putatively a novel *Gemmatimonadetes* or *Firmicutes*, also had annotations for phototrophy.

Lastly, bins annotated as *Bacteroiodetes* were the most common in the study (bins 5,11,12,13,14,15,17,18) and contained annotations for aerobic heterotrophy, represented by cytochrome c oxidase (CCON), glycolysis genes enloase (ENO), pyruvate kinase (PYK), and 6-phosphoglucanate dehydrogenase (PG). Despite their multitude, functional differences between these bins were not identifiable using our biogeochemical cycling gene analysis.

### Sequence variation analysis of *C. chthonoplastes* PCC 7420 by coverage, subsystems, and genes

The poor assembly statistics for the dominant bin identified in these mats (bin 1; most similar to *C. chthonoplastes*) suggested that genetic subpopulations of this organism confounded the assembler. We used the GenBank draft genome of the type strain PCC 7420, isolated from hypersaline microbial mats, as a mapping scaffold for metagenome reads. All variants (SNPs, indels, repeats) were inferred, but only SNPs, which were the most common variant, were retained for downstream analysis. SNPs occurred in lower coverage regions compared to the coverage distribution of the sequenced genome (Figure 4). These variants represent a rarer strain, or strains, with ~150x coverage in the sample while the main coverage distribution at ~600x coverage had fewer SNPs. SNPs from these lower coverage regions (50-200x coverage) were filtered and summarized for each gene and each subsystem. The difference between the variant density of a subsystem to the genome average was tallied for all genes in a subsystem to form a score to evaluate the relative accumulation of mutations across gene subsystems. This score weighted for subsystems with more genes. Subsystems with less variant accumulation had a negative score, and subsystems with more variant accumulation had a positive score (Supplemental Figure S5). Scores from subsystems that had the least and most accumulation are shown in Table 2. Subsystems with low variant density scores were related to photosynthesis (e.g. photosystem I/II, phycobilisome, chlorophyll), carbon fixation (e.g. cAMP, carboxysome, circadian clock), and basic cellular processes (e.g. cell division, DNA replication, ribosomal proteins, respiration, central metabolism). Examining individual genes from these subsystems with high variant density, specific genes that might indicate strain differentiation (Table 3) were regulatory proteins (P-II, kinases, cAMP proteins), membrane proteins (Co/Zn/Cd efflux, phosphate permease, O-antigen export permease, isoprenoid and carotenoid biosynthesis, fatty acid biosynthesis, and amino-sugar biosynthesis), nitrogen and amino acid cycling genes, and several transferases, and many genes related to carbohydrate modification. Additionally, environmental stress response genes, represented by chemotaxis genes, cryptochrome, and Exodeoxyribonuclease, and DNA-cytosine methyltransferase, were observed. Supplemental Dataset 2 contains the full list of subsystem and gene variant density statistics.

**Figure 4.**
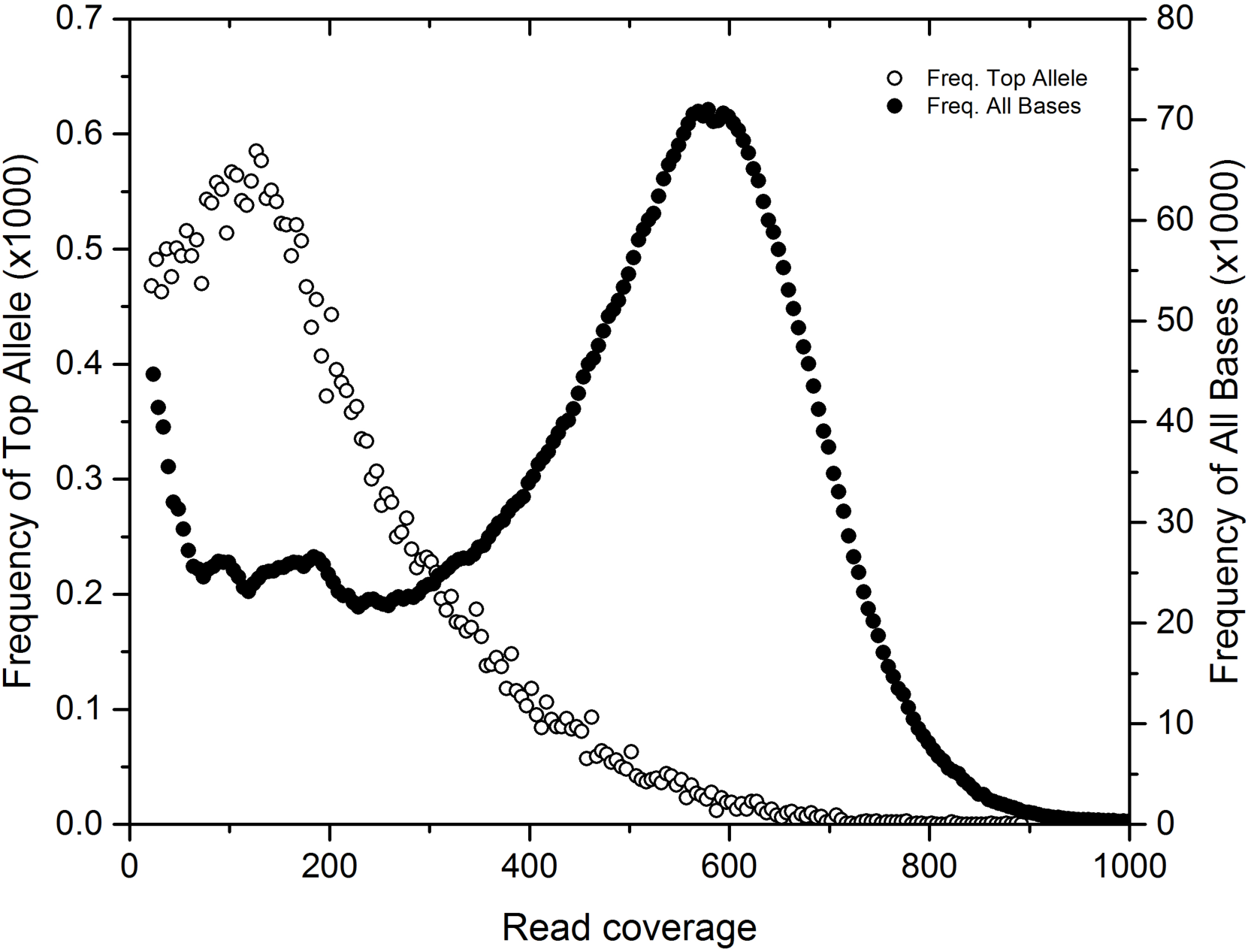
PCC7420 SNP coverage mapping overview showing total base coverage (closed circles, right axis) and found SNP dominant allele coverage (open circles, left axis) indicating SNP allele prevalence at lower coverage.

**Table 2.** SNP alleles in subsystems indicating variable and conserved gene categories

|  | Number of genes between variant rates of (upper inclusive): |  |  |  |  | Score |
| --- | --- | --- | --- | --- | --- | --- |
|  | all | 0-1% | 1-2% | 2-3% | +3% |  |
| Most conserved subsystems |  |  |  |  |  |  |
| cAMP signaling in bacteria | 87 | 52 | 11 | 0 | 0 | -24.2 |
| CO2 uptake, carboxysome | 64 | 35 | 10 | 0 | 0 | -15.6 |
| Photosystem II | 21 | 4 | 0 | 0 | 0 | -8.9 |
| Bacterial Cell Division | 25 | 14 | 2 | 0 | 0 | -8.3 |
| Phycobilisome | 18 | 2 | 0 | 0 | 0 | -7.7 |
| DNA-replication | 26 | 16 | 3 | 0 | 0 | -7.6 |
| Entner-Doudoroff Pathway | 20 | 9 | 1 | 0 | 0 | -7.3 |
| Ribosome SSU bacterial | 16 | 2 | 0 | 0 | 0 | -6.8 |
| SigmaB stress responce regulation | 23 | 8 | 3 | 0 | 0 | -6.3 |
| Chlorophyll Biosynthesis | 17 | 10 | 1 | 0 | 0 | -6.1 |
| Respiratory Complex I | 14 | 6 | 0 | 0 | 0 | -6.0 |
| Cyanobacterial Circadian Clock | 35 | 21 | 4 | 1 | 0 | -5.8 |
| Peptidoglycan Biosynthesis | 19 | 14 | 2 | 0 | 0 | -5.8 |
| Photosystem I | 12 | 2 | 0 | 0 | 0 | -5.1 |
| Ton and Tol transport systems | 25 | 16 | 1 | 1 | 0 | -5.1 |
| Most variable subsystems |  |  |  |  |  |  |
| Fatty Acid Biosynthesis FASII | 27 | 7 | 3 | 3 | 1 | 20.2 |
| CBSS-258594.1.peg.3339 (glycotransferases) | 35 | 16 | 2 | 4 | 1 | 20.1 |
| Polyprenyl Diphosphate Biosynthesis | 5 | 2 | 0 | 1 | 1 | 17.2 |
| Isoprenoid Biosynthesis | 11 | 3 | 1 | 1 | 1 | 15.8 |
| DNA repair, bacterial | 23 | 12 | 1 | 2 | 1 | 15.1 |
| Cobalt-zinc-cadmium resistance | 10 | 5 | 3 | 0 | 1 | 14.2 |
| Pentose phosphate pathway | 4 | 3 | 0 | 0 | 1 | 13.2 |
| Carotenoids | 15 | 10 | 0 | 1 | 1 | 13.0 |
| Rhamnose containing glycans | 12 | 3 | 4 | 3 | 0 | 12.8 |
| Bacterial Chemotaxis | 26 | 17 | 3 | 1 | 1 | 11.8 |
| Glutamine, Glutamate, Aspartate and Asparagine Biosynthesis | 13 | 8 | 2 | 0 | 1 | 11.7 |
| Ammonia assimilation | 12 | 5 | 1 | 0 | 1 | 11.0 |
| Bacterial RNA-metabolizing Zn-dependent hydrolases | 10 | 6 | 0 | 0 | 1 | 10.7 |
| High affinity phosphate transporter and control of PHO regulon | 13 | 5 | 1 | 3 | 0 | 8.9 |
| Conserved gene cluster associated with Met-tRNA formyltransferase | 17 | 13 | 1 | 0 | 1 | 8.9 |
| Maltose and Maltodextrin Utilization | 14 | 5 | 5 | 2 | 0 | 8.7 |
| Glutathione: Biosynthesis and gamma-glutamyl cycle | 5 | 2 | 1 | 2 | 0 | 7.9 |
| Sialic Acid Metabolism | 5 | 1 | 1 | 2 | 0 | 7.9 |
| Calvin-Benson cycle | 17 | 6 | 0 | 0 | 1 | 7.7 |
| Average |  |  |  |  |  | 0.0 |
| Median |  |  |  |  |  | -1.4 |
| Standard Deviation |  |  |  |  |  | 5.8 |

## Discussion

### Metagenomic binning can reproduce the systems biology of microbial mats by providing a genetic atlas

In this study, *de novo* metagenomic binning approaches were used to reconstruct the biogeochemical cycling divisions between different organisms of a microbial mat and to seek novel diversity previously unrevealed by reference-based annotation studies. Phylum level metagenomic results concur with past findings of the overall types of *Proteobacteria, Cyanobacteria,* and *Bacteroidetes* observed in Elkhorn Slough mats (Burow et al. 2012) and with previous metagenomic studies on lithifying and non-lithifying hypersaline mats (Khodadad & Foster 2012; Harris et al. 2013; Ruvindy et al. 2016). Bins 1 and 2 were annotated as the filamentous *Cyanobacteria* typically observed in mat systems, i.e. the genera *Coleofasciculus* and *Lyngbya*, respectively. Bin 3 was annotated as a sulfide-oxidizing bacteria from a *Thiorhodovibrio* sp., which are commonly observed in mats (Overmann et al. 1992). Bin 10 supported observations of nitrogen fixation potential in purple nonsulfur bacteria in mats (Bebout et al. 1993; Zehr et al. 1995) but also suggests the multifaceted role (i.e. sulfate and nitrate metabolism) of many PNS bacteria in microbial mats (Yurkov et al. 1994). Observations of sulfate reduction in the phototrophic zone of mats (Canfield & Des Marais 1991; Fike et al. 2008) was supported by one *Deltaproteobacteria* bin (bin 16). This bin, with closest relative *Desulfotalea psychrophila* LSv54, contained annotations for sulfate reduction pathways and a methyl viologen-reducing hydrogenase suggested as markers for SRBs by previous work (Pereira et al. 2011; Burow et al. 2014). This bin also contained a number of annotations that indicate oxygen tolerance and support previous studies that identified sulfate reduction in the phototrophic zone of hypersaline microbial mats and suggested that oxygen-tolerant SRBs were responsible (Canfield & Des Marais 1991; Visscher et al. 1992; Jørgensen 1994; Teske et al. 1998; Baumgartner et al. 2006; Fike et al. 2008; Burow et al. 2014). However, isolation of such organisms has been difficult and multiple mechanisms of oxygen tolerance have been proposed (Dolla et al. 2006). Similar to previous findings, the bin we identified does not appear to contain cytochrome c oxidase or superoxide dismutase (Burow et al. 2014) but does have rubrerythrin, suggesting that this organism may scavenge oxygen using this pathway or have a syntrophic oxygen coping strategy seen in mat-derived SRBs (Sigalevich et al. 2000).

Finally, when examining organoheterotrophs in this subset, numerous bins corresponding to the phylum *Bacteroidetes* were found. Heterotrophic diversification in the top 2 mm of Elkhorn Slough mats appears to be prevalent, likely because these systems are replete with fixed carbon derived from spent fermentation products (Lee et al. 2014), exuded polysaccharides from *Cyanobacteria* (Stuart et al. 2015), and potentially other unobserved carbon influxes such as agricultural and animal inputs. A similar phenomena is observed in saccharide-rich gut ecosystems (Eckburg et al. 2005; Gill et al. 2006) where abundant carbohydrate substrates have been linked to great metabolic variety among *Bacteroidetes* at the species and even strain level (Cottrell & Kirchman 2000, Rogers et al. 2013). Additional emphasis on the heterotrophic niche partitioning of microbial mat ecosystems is needed to reveal which major factors such as light cycle, substrate variety, and electron donors drive heterotrophic diversification.

We noted that not all canonical members of microbial mats were observed, even when unbinned sequences were searched. Notably missing from this study were the chemolithotrophic sulfur bacteria (*Beggiotoa* sp.) and methanogenic Archaea, both of which have been observed from Elkhorn Slough microbial mats (unpublished data). Deeper sequencing efforts may be required to detect enough genomic sequence to bin these rarer clades, as was also noted for ESFC-1; alternatively sampling from deeper mat depths may be required to capture these additional species.

### *C. chthonoplastes* is distinguished from other *Cyanobacteria* in mats by extensive polysaccharide production and breakdown capability

To better understand *C. chthonoplastes* ubiquity in microbial mats, we compared the three *Cyanobacteria* (bins 1,2,9) plus the genome of filamentous cyanobacterium ESFC-1 (isolated previously from this site but poorly assembled in this dataset) for major metabolic differences, specifically for carbon utilization genes (Figure 2). Notably, the dominant bin (*C. chthonoplastes*) appears to have an extensive capability to synthesize and process beta-glucoside polymers, making it distinct in this study and supporting recent work showing that fixed carbon polysaccharide production by *Cyanobacteria* (notably beta-glucose polysaccharides) constitutes a sizable fraction of extracellular polysaccharide (Stuart et al. 2015). The elevated mutation levels seen in these genes also support the view that polysaccharide regulation plays a role in strain differentiation in this species. The role of these beta-glucose polymers are not well understood, but may play a role in carbon storage for *Cyanobacteria* (Stuart et al., 2015), or may have adhesion and anchoring functions (Ross et al. 1991; Römling 2002). These two details infer a phenotype of polysaccharide specialization that both distinguishes the dominant mat-building *Cyanobacteria* found in these mats and differentiates strains of this species from each other.

### Metagenomic binning predicts novel functional roles of microbes from microbial mats

Binning was used to identify novel organisms from unstudied clades to examine the novel diversity in microbial mats. These binning efforts revealed that *Gammaproteobacteria* consisted not only of the canonical sulfide-oxidizing bacteria in mats, but also of bins with bacteriochlorophyll-containing heterotrophs, possibly containing mixotrophs similar to OM60 clades (Spring & Riedel 2013). As OM60 clades have been shown to rely on specific light, oxygen, and organic acids for mixtotrophic growth, this work suggests that the isolation of such bacteria would be highly dependent on mimicking specific conditions that develop during a diel cycle, such as the acetate-replete, low-oxygen, low-light initial early morning photosynthetic period of microbial mats, where their growth would be distinctive from other heterotrophs. At present, given the lack of genomic knowledge about phototrophic *Gammaproteobacteria*, accurate taxonomic assignment of these bins below the class level remains challenging. This work also identifies a potential novel phototroph within *Firmicutes* or *Gemmatimonadetes.* Based on the taxonomic identity and binning completeness we measured, our research suggests this bin represents a possible salt-tolerant variant of the recently isolated phototrophic *Gemmatimonadetes*. Sequences of taxa from freshwater lakes belonging to possible clades of phototrophic *Gemmatimonadetes* (Zeng et al. 2014) bear some resemblance to this bin. Specifically, the annotations of photosystem genes in this bin appear to be derived from *Alphaproteobacteria*, and contain annotations for aerobic respiration genes (cytochrome c oxidase (CCO), (2-oxoglutarate synthase) KORA, enolase (ENO), pyruvate kinase (PYK), and 6-phosphoglucanate dehydrogenase (PGD), Figure 3). However, given the possibility of misassembly, mis-annotation, and the difficulty of inferring gene expression from genomic data, these findings must be paired with further microbial isolation, characterization, and genome sequencing.

### Reference-based variant analysis reveals core and accessory genomes *C. chthonoplastes*

Our analyses indicate that genomic flexibility of subpopulations of *C. chthonoplastes* can be detected by metagenomic variant analysis. Comparative genomic analysis has shown that species can contain recombinant and horizontally transferred regions that allow for both core and flexible genome elements (Fraser et al. 2007; Kashtan et al. 2014). Differential coverage analysis suggested that the dominant strain resembled the sequenced PCC 7420, but that at least one more rare subtype with less similarity was also present (Figure 4). Focusing on these less abundant variants, each SEED subsystem was assessed for the density of variant sites in gene subsystems. In subsystems related to carbon metabolism, phototrophy, DNA replication, and cell division we observed reduced levels of variant accumulation that seems consistent for the essential functioning subsystems of a cyanobacterium (Table 2). This suggested that assessment of variability can predict conserved subsystems that represent the core genome of that organism. Conversely, subsystems associated with greater accumulation of mutations should indicate potential variation within niches. A metagenomic study of meltwater mats from Arctic and Antarctic ice shelves (Varin et al. 2012) suggested that coping with environmental regulation, especially in variable salinity conditions, was a primary driver of genetic functional diversity. Similarly, in this study, subsystems displaying increased variation suggest that C, N, P nutrient cycling and environmental stress response to factors such as salinity, metals, light and infection were the drivers in *C. chthonoplastes* genetic differentiation in mats. Specific genes that had increased variation (Table 3) included a nitrogen regulatory gene involved in modulating nitrogen scavenging, a transaldolase (part of the pentose-phosphate pathway) and several glycotransferases and carbohydrate processing genes, an endonuclease involved in DNA repair, a DNA methyl-transferase related to phage immune response, a metal ion pump involved in toxicity resistance, and a number of fatty acid, carotenoid, isoprenoid biosynthesis pathway genes. This work predicts that these factors can be used to differentiate *C. chthonoplastes* strains, both physiologically and genetically, and may explain some of the ubiquity of this species across microbial mats worldwide.

**Table 3.**
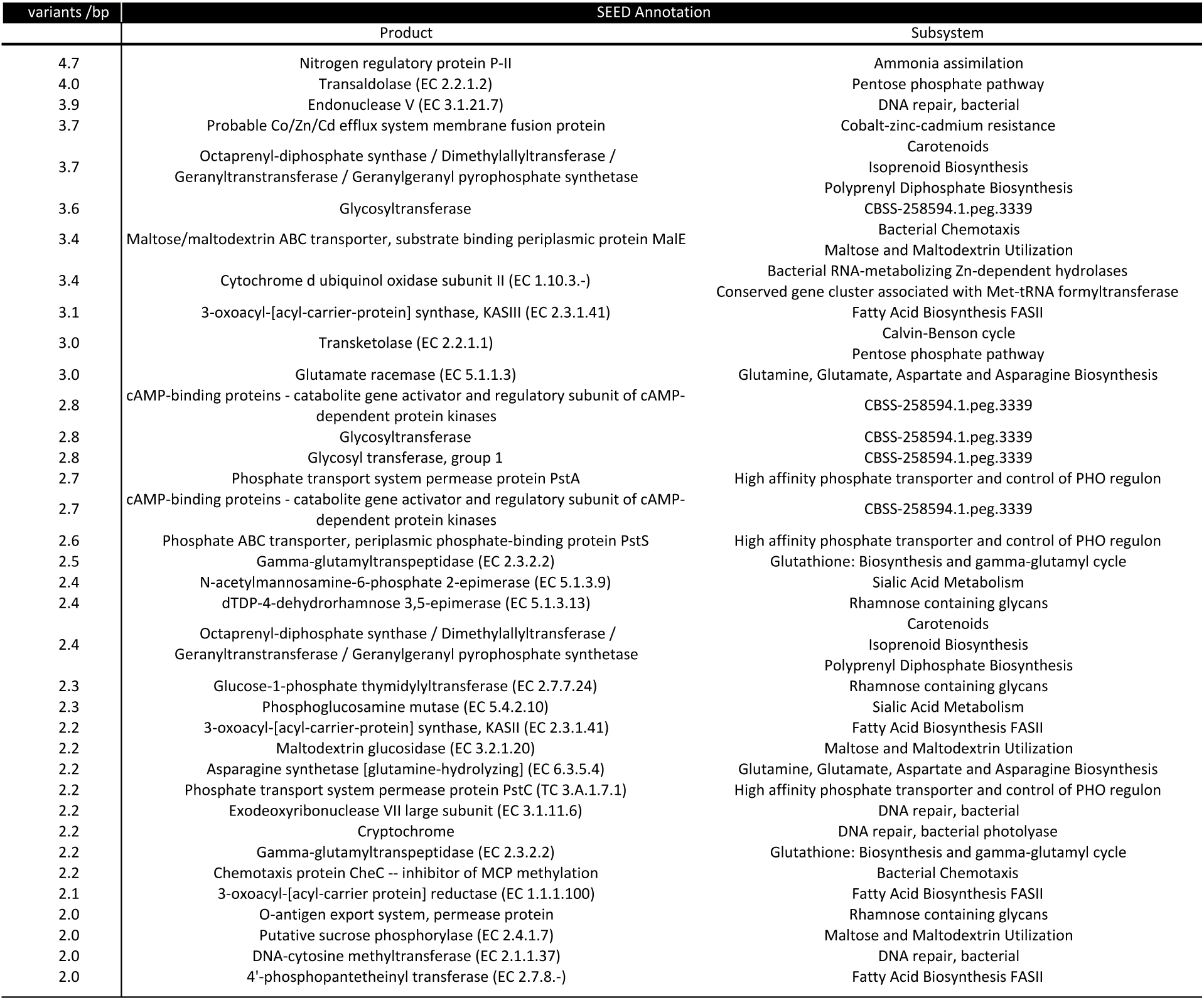
Variant density (expressed as variants per gene length) for individual genes from subsystems with high variance score (excluding unknown genes and genes from unknown categories).

## Conclusion

In this study coverage binning was used to resolve functional genes at a genome population level and thus create a genetic basis for biogeochemical partitioning that parallels results seen from physiological and biogeochemical cycling studies. The use of reference-free binning was crucial as the majority of bins identified had only a fraction of genes matching any nearest reference genome. We also show that where reference genomes were available, a key weakness of assembly-based analysis, strain level microheterogeneity, can be used to generate SNP analysis to differentiate strain level differences using metagenome reads. Though the ecosystem survey in this study included a limited snapshot of a microbial mat, using only abundant organisms and key genes in metabolic pathways, we can see how the ecosystem roles of mat microbes partition the genetics within metagenomes. Our analysis also predicts a number of novel mat organisms that are as yet unidentified, including several phototrophs. We suggest that these combined reference and reference-free analysis approaches can be used to generate genetic atlases of biogeochemical cycling of novel ecosystems and direct further microbial investigation.

## Acknowledgements

This project is funded by the Department of Energy through the Genome Sciences Program under contract SCW1039 to the Lawrence Livermore National Laboratory (LLNL) Biofuels Scientific Focus Area and performed under the auspices of LLNL under Contract DE-AC52-07NA27344. This research used resources of the National Energy Research Scientific Computing Center (NERSC) and the DOE Joint Genome Institute (JGI) Community Sequencing Program award #701; both DOE Office of Science User Facilities are supported by the Office of Science of the U.S. Department of Energy under Contract No. DE-AC02-05CH11231. RCE acknowledges the support of a NASA Postodoctoral Fellowship. The authors wish to thank Rob Egan at NERSC for assistance with Ray-Meta, Tijana Glavina del Rio at JGI for sequencing assistance, and Luke Burow, Mike Kubo, and Tori Hoehler for assistance with diel sampling. We thank Jeff Cann, Associate Wildlife Biologist, Central Region, California Department of Fish and Wildlife, for coordinating our access to the Moss Landing Wildlife Area.

## Conflicts of Interest

Authors declare no conflict of interest

## Supplemental Figures

**Figure S1:**
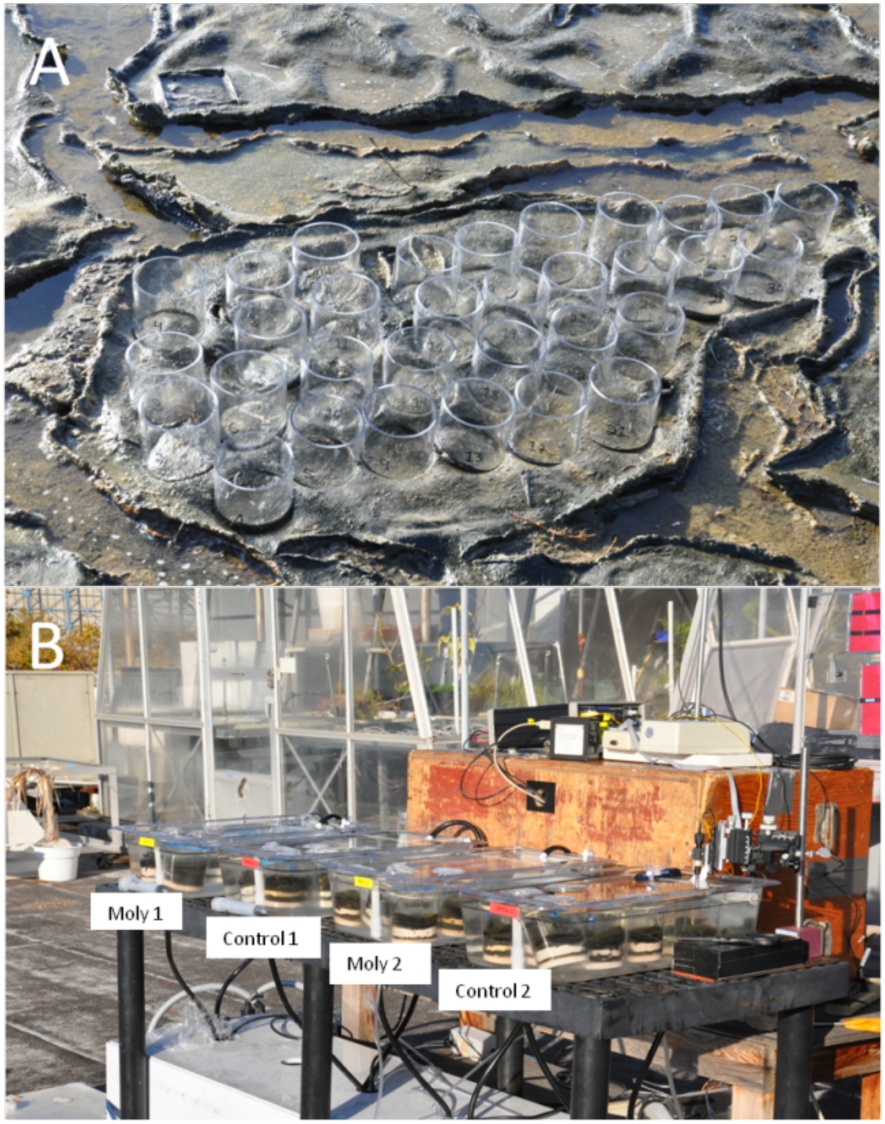
A. Photograph of location of cores collected in the field from the microbial mats in Elkhorn Slough, Moss Landing, California on November 7, 2011. Individual samples collected in core tubes were numbered and could be tracked throughout the diel experiment. B. Experimental apparatus used to incubate microbial mats throughout the diel period. Incubation containers containing cores used for control and molybdate treatments are labeled.

**Figure S2:**
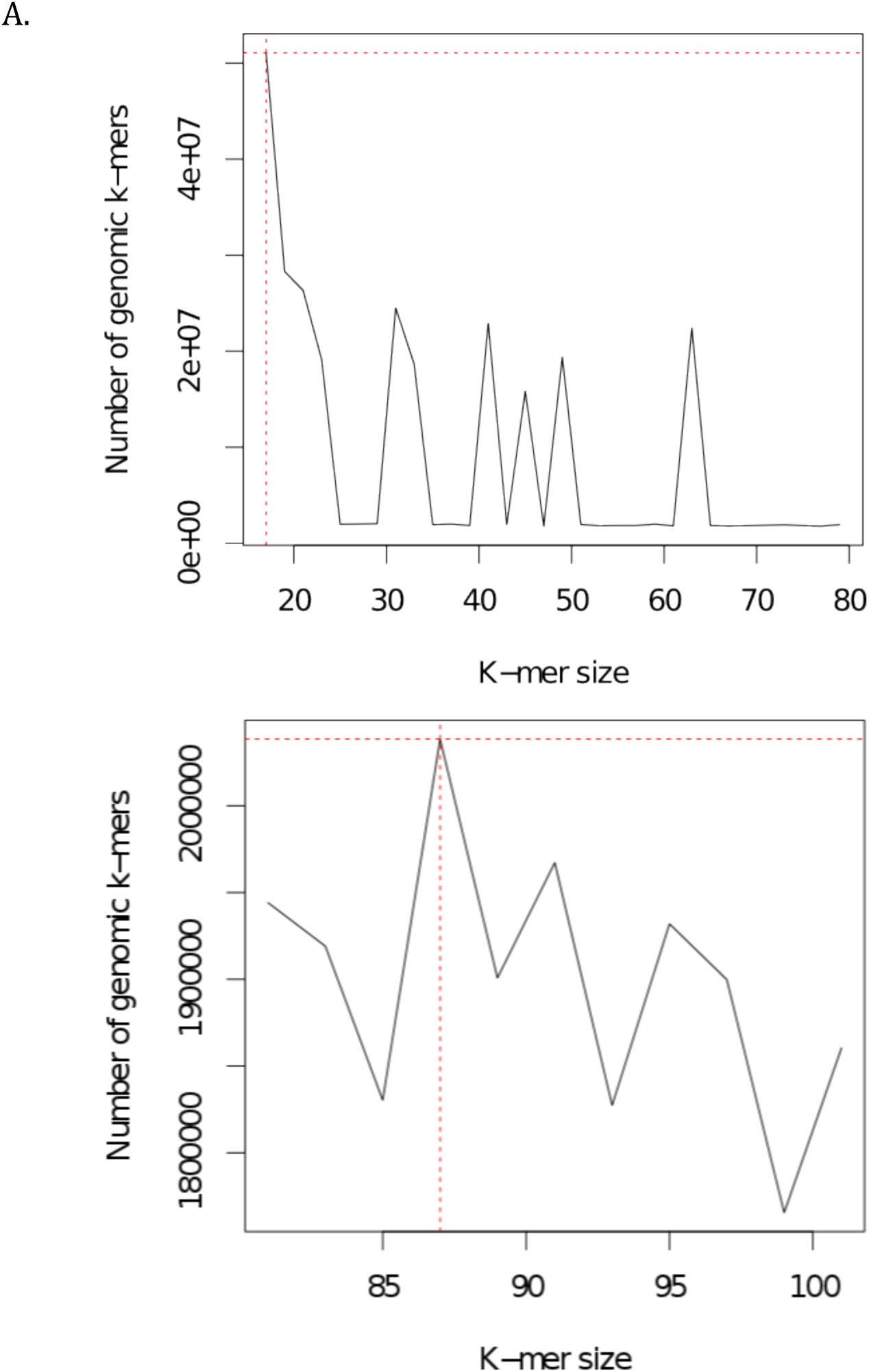
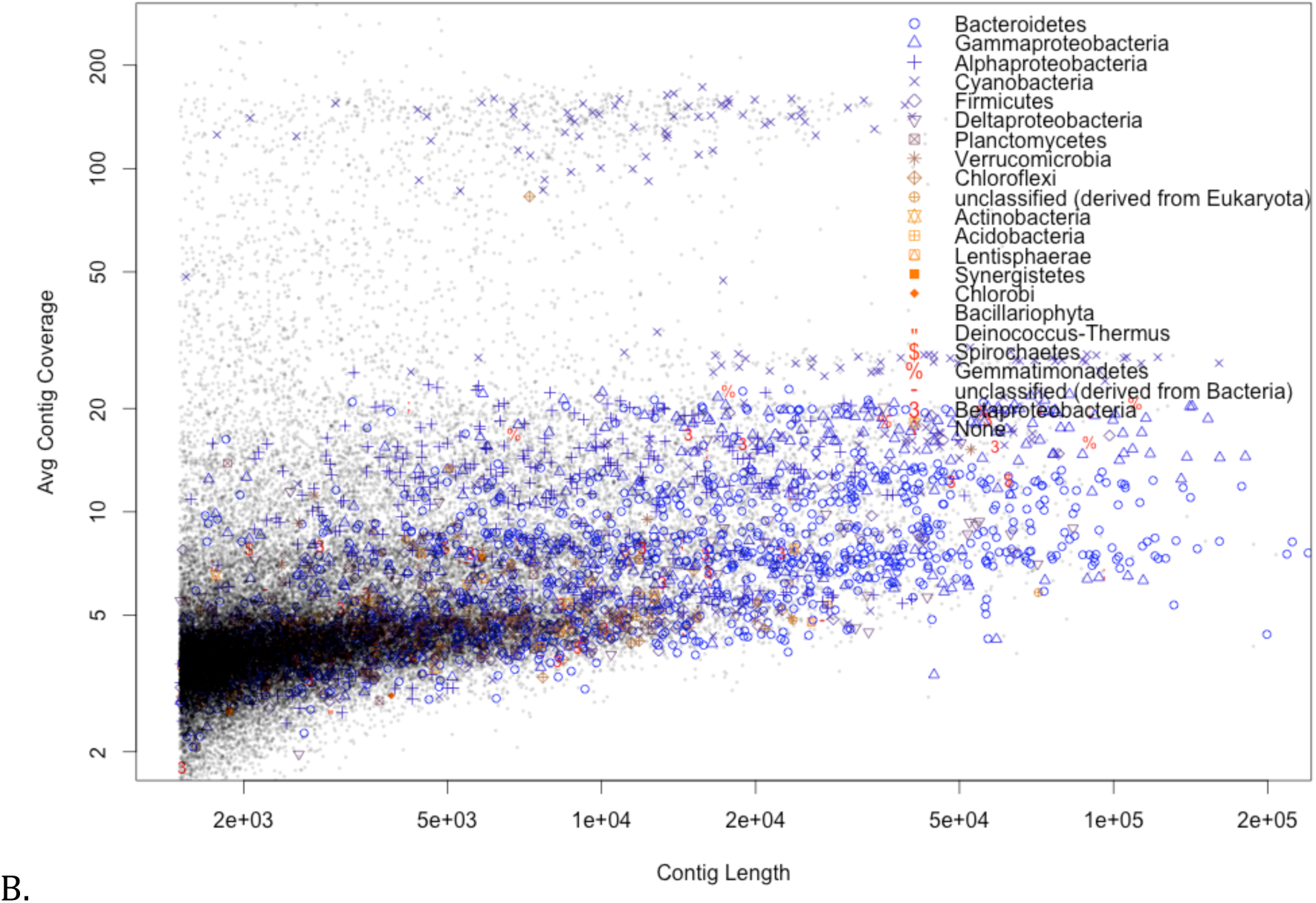
Kmergenie plots indicating optimal word sizes for assembly (k=29,45,63 were chosen) (A) and coverage v. length for k=29 word assembled scaffolds indicating a phylogenetic signal relating to contig size and coverage (B).

**Figure S3:**
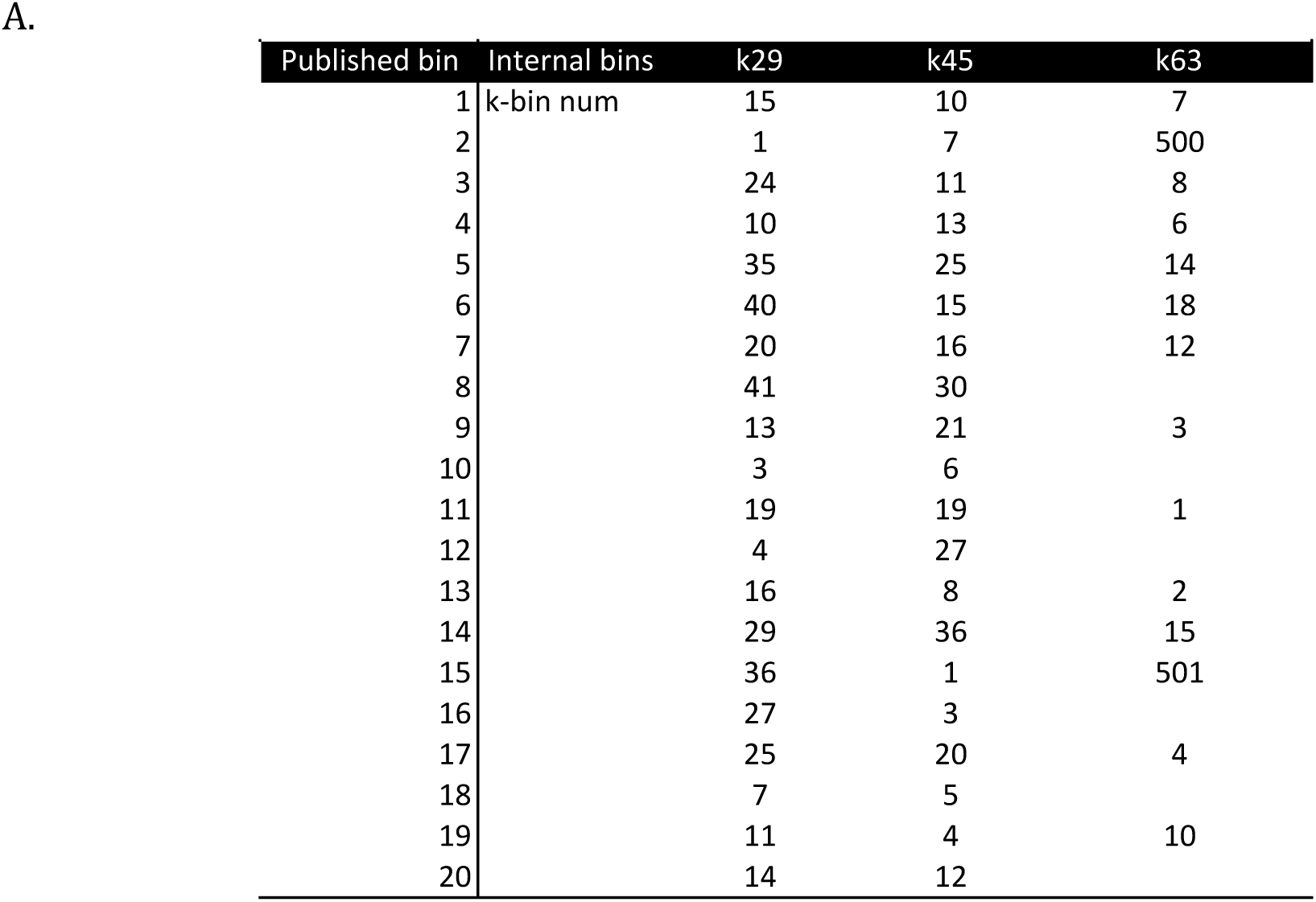
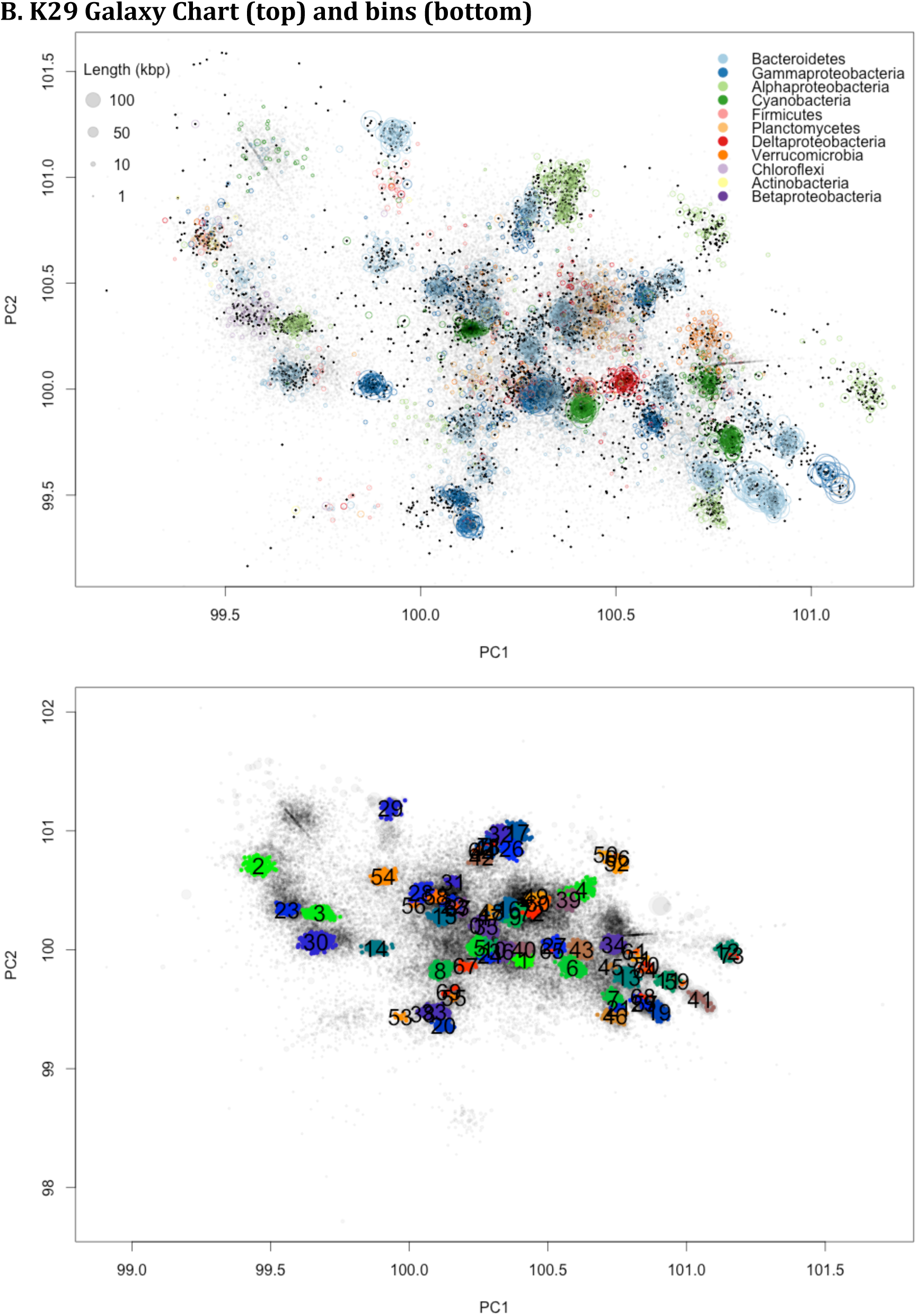
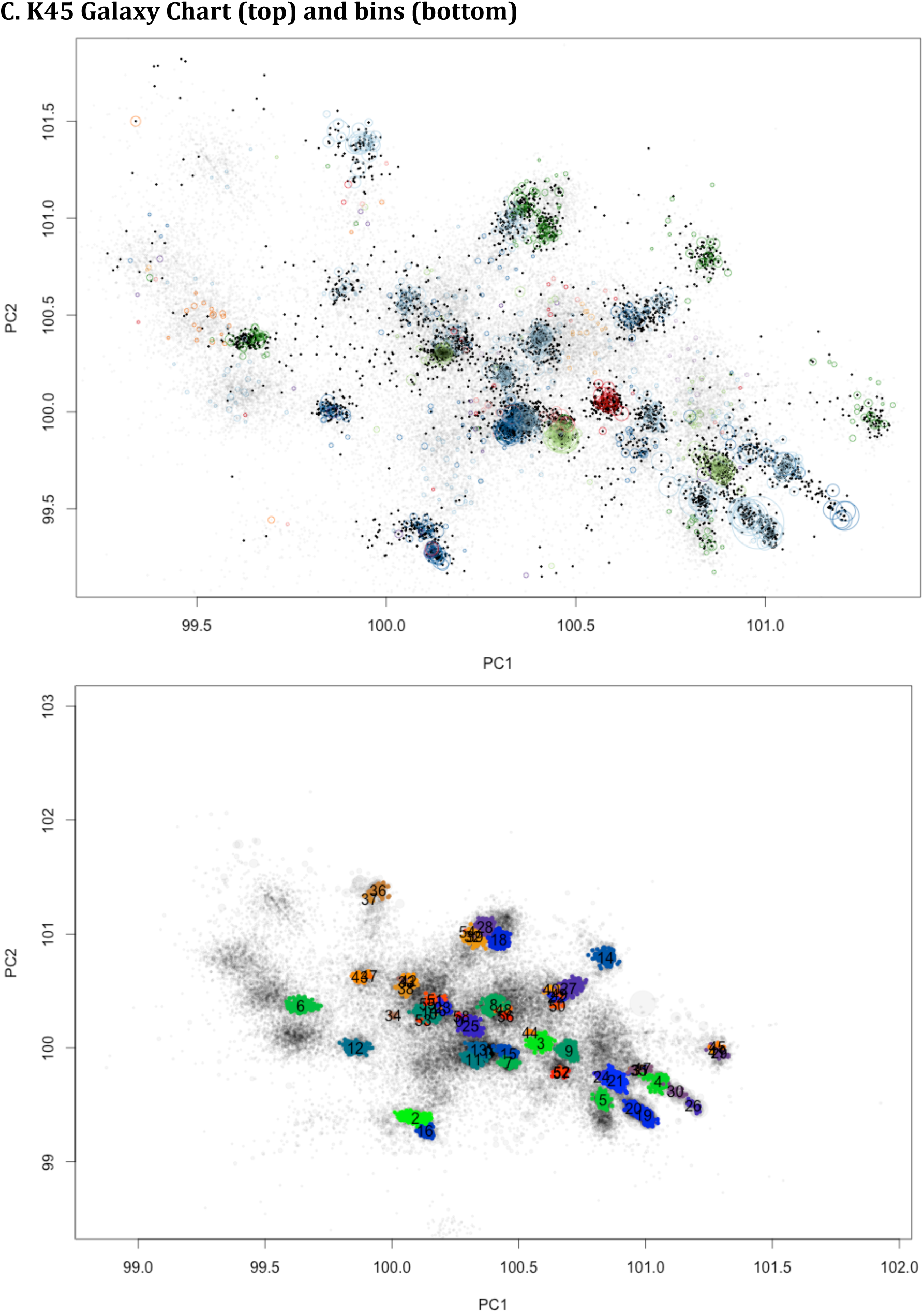
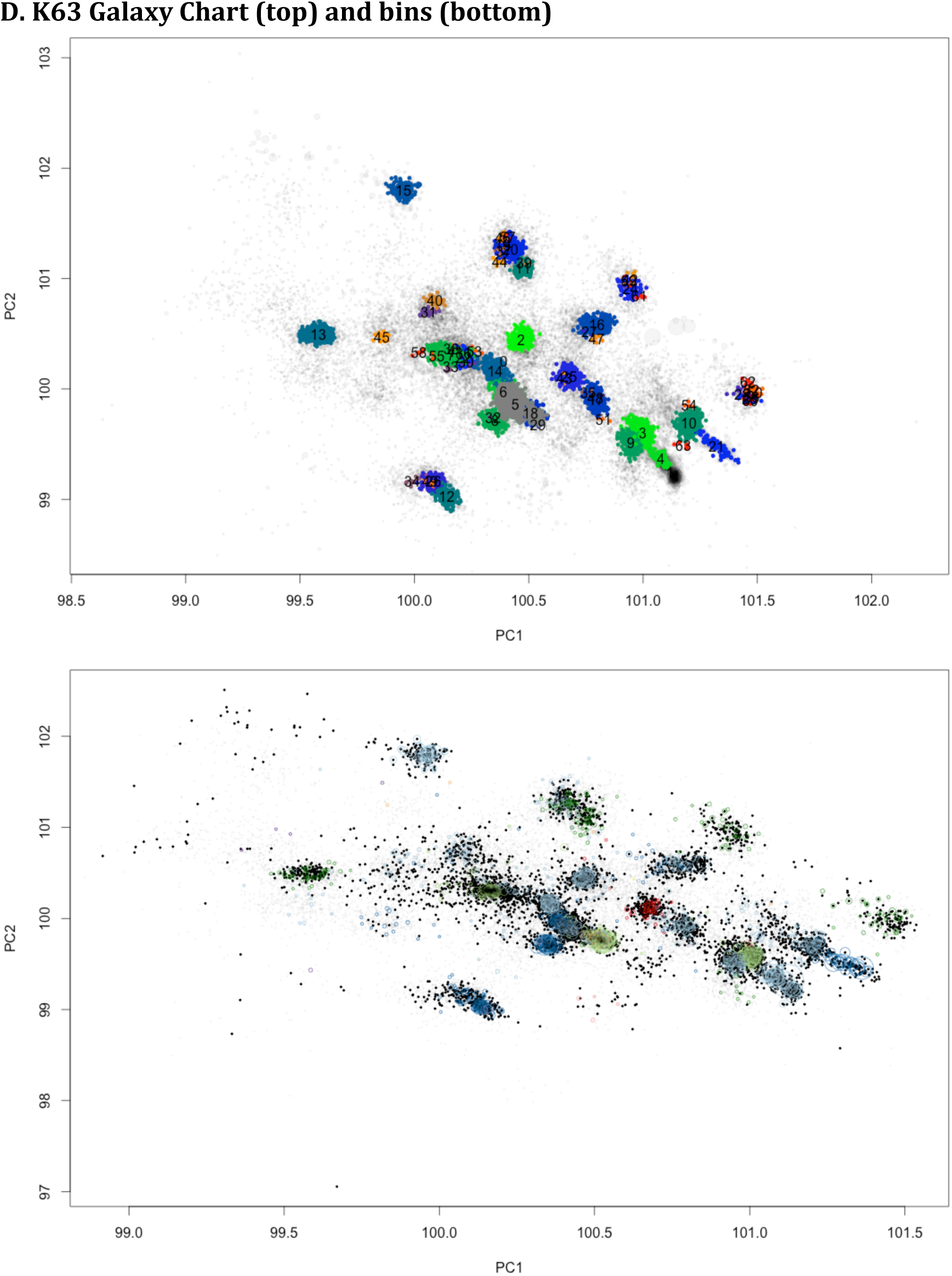
Bin reference table (A), and k=29,45,63 word size PCA galaxy charts and bin charts (B, C, D)

**Figure S4:**
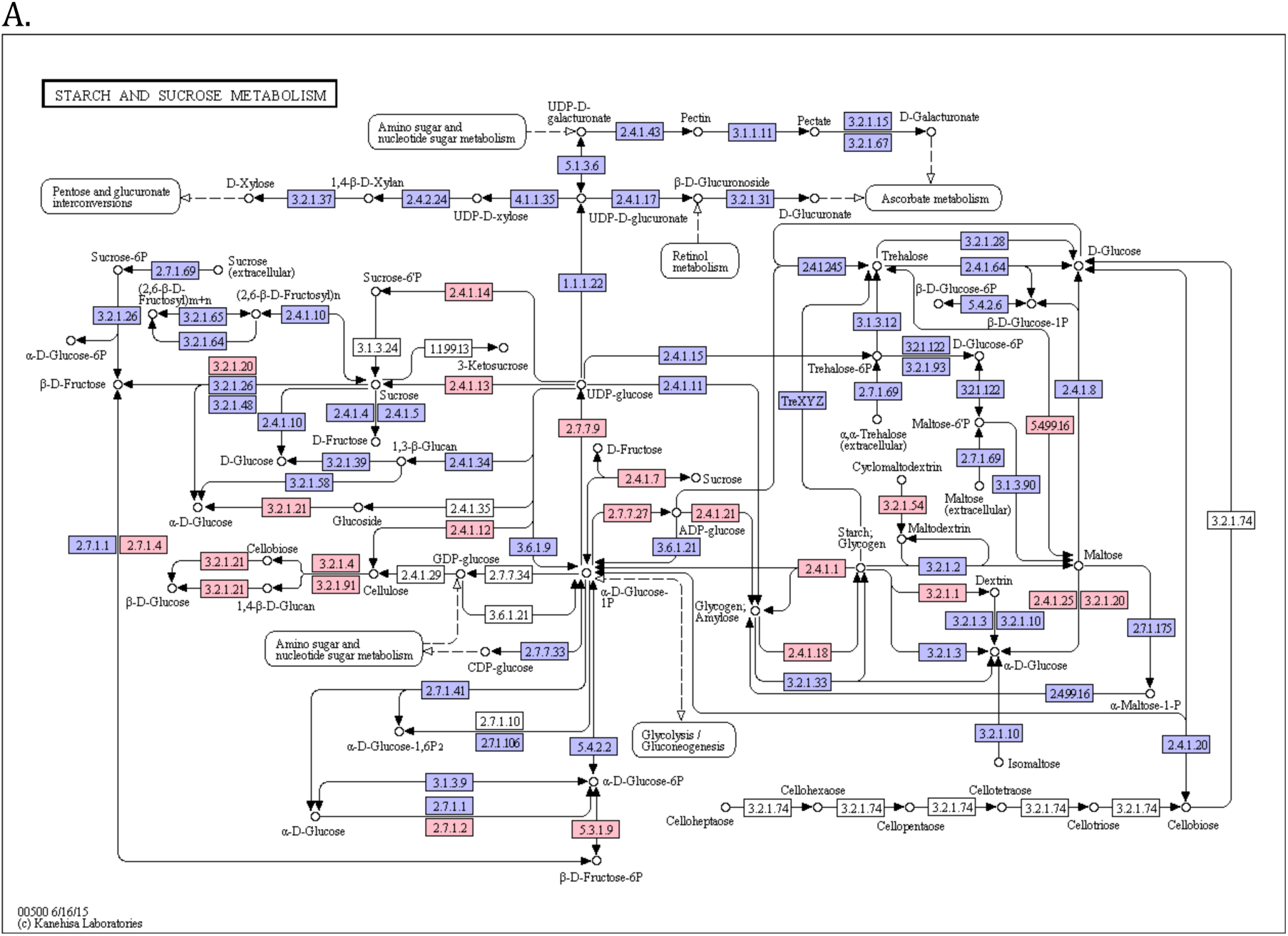
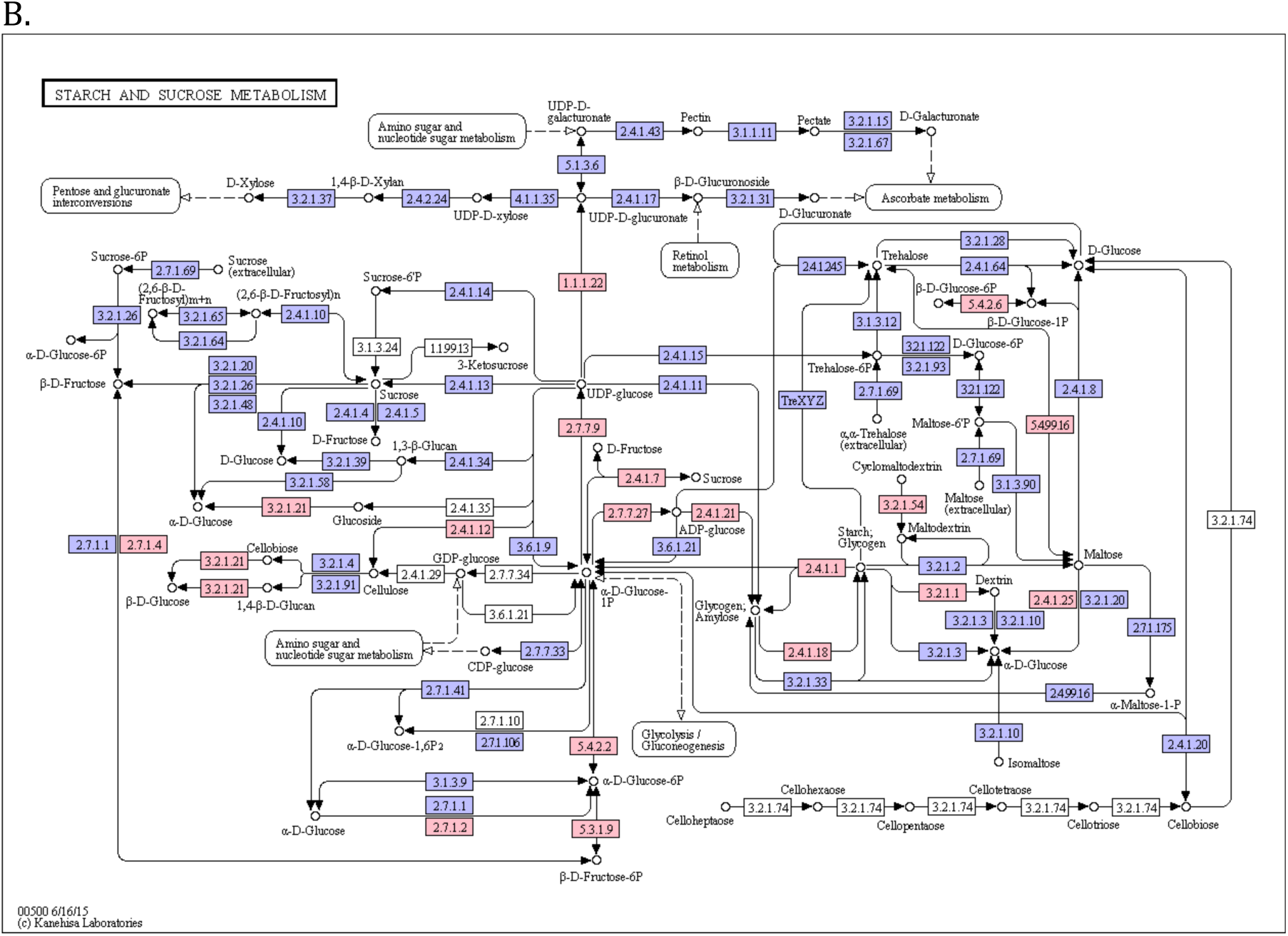
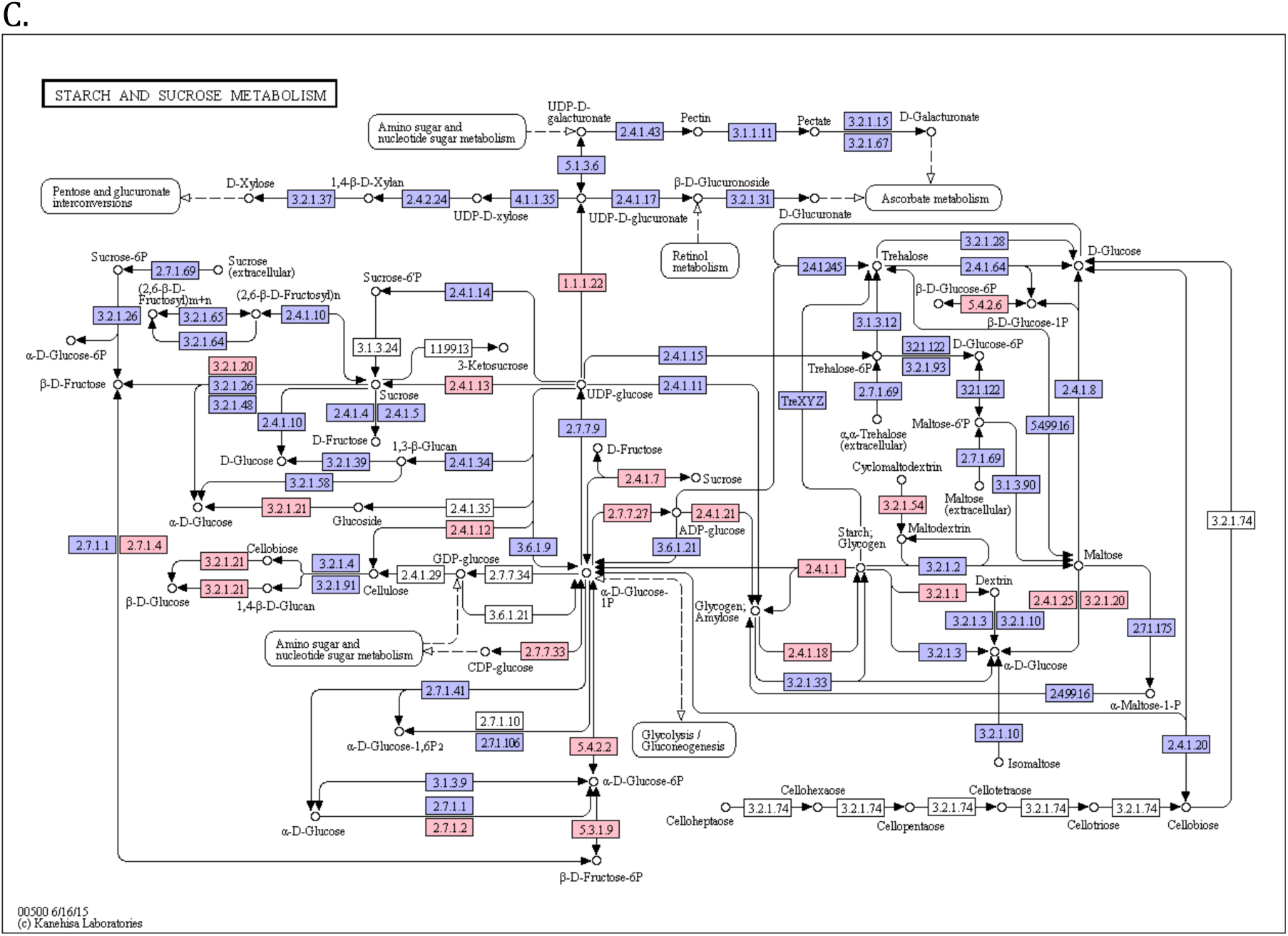
Preliminary KEGG figures of starch and sucrose metabolic pathways in each Cyanobacteria-annotated bin based on MG-RAST KEGG annotations (red indicates present, blue indicates absent). k63.7, bin 1, Coleofaciculus chthonoplastes PCC7420 (A), k29.1, bin 2, Lyngbya spp. (B), k29.13, bin 9, unknown Cyanobacterium (C).

**Figure S5:**
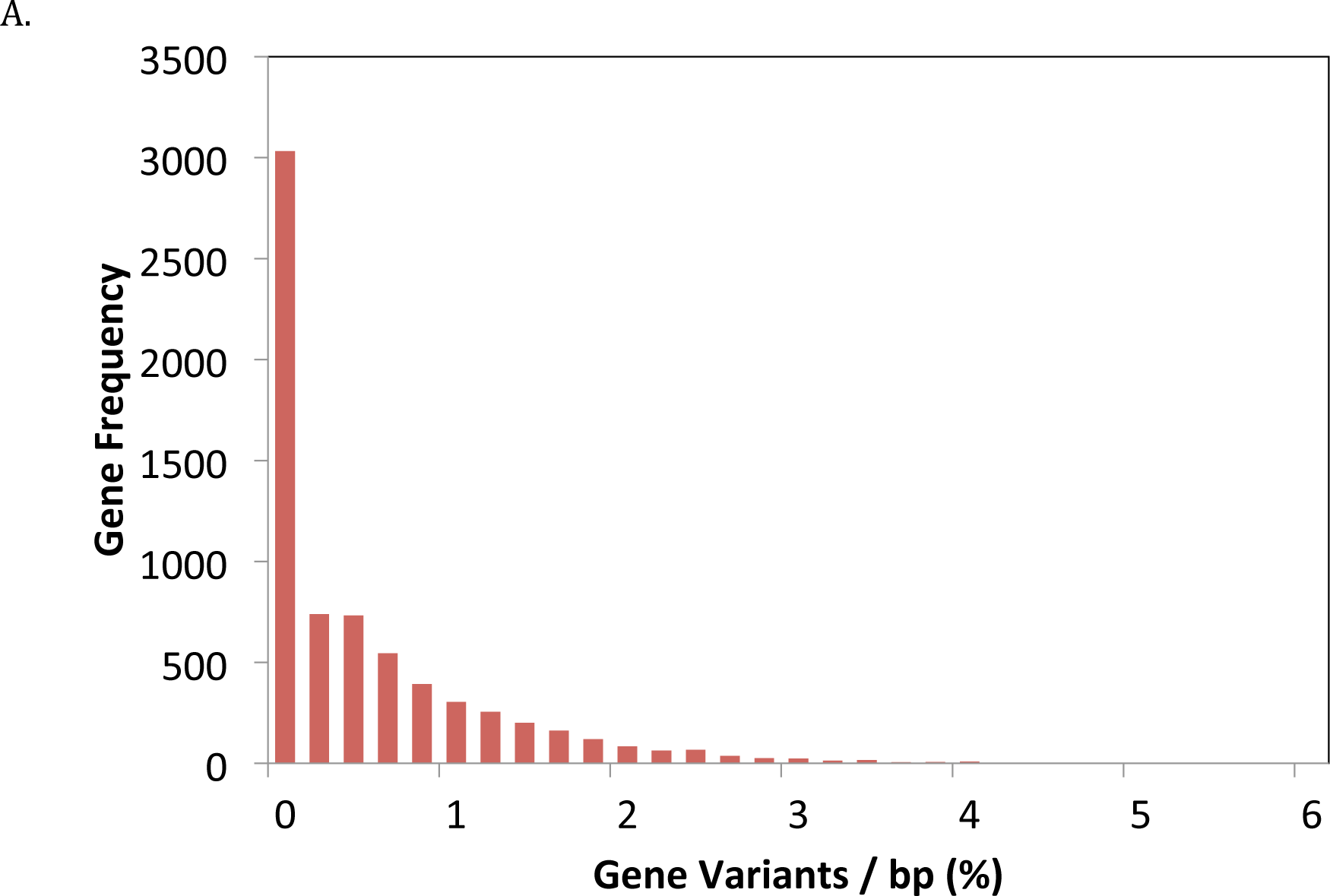
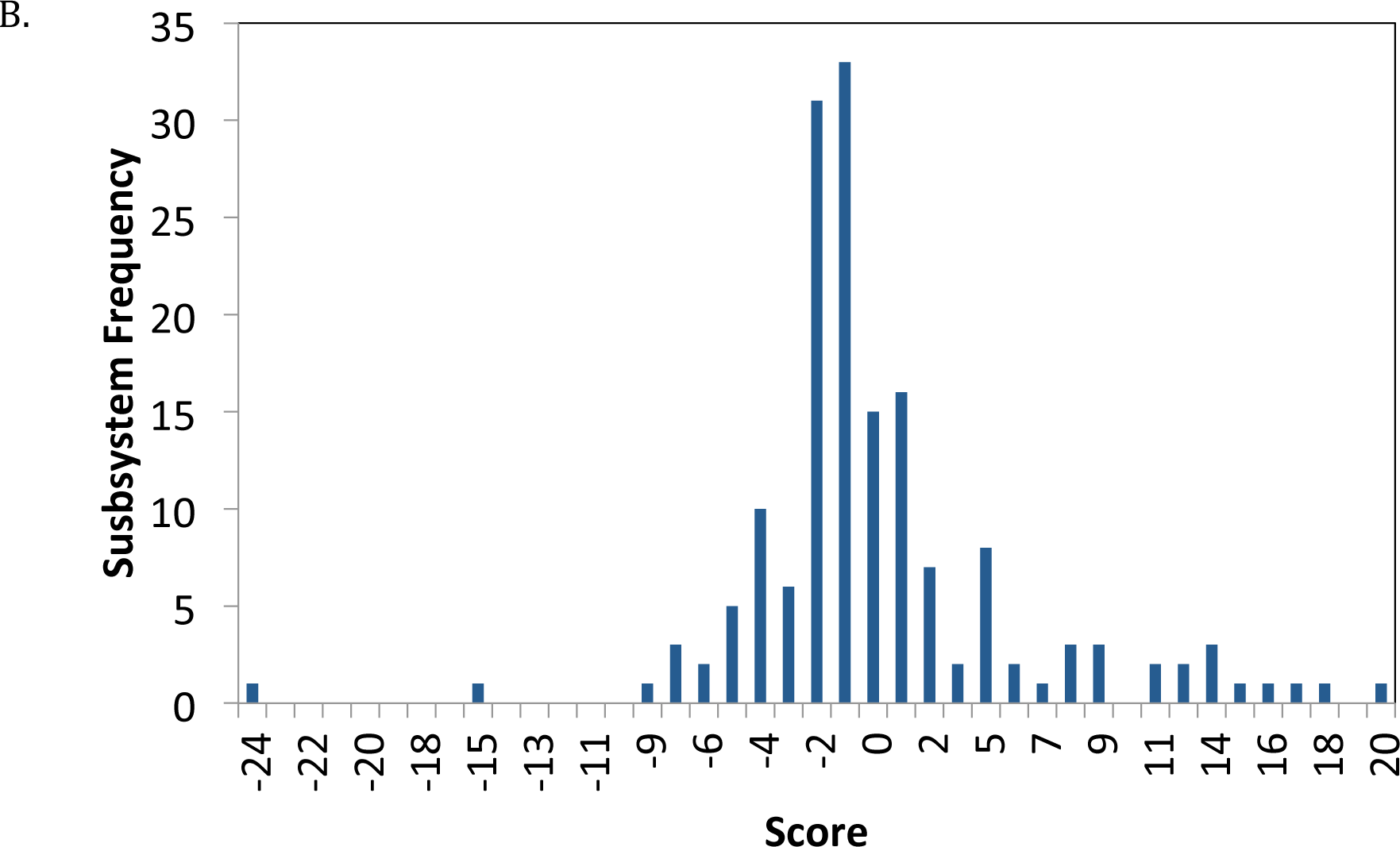
Histogram of variants / bp of all genes in PCC 7420 (A), and histogram of subsystems scores (B)

## References

Albertsen M, Hugenholtz P, Skarshewski A, Nielsen KL, Tyson GW, Nielsen PH. (2013). Genome sequences of rare, uncultured bacteria obtained by differential coverage binning of multiple metagenomes. Nat. Biotechnol. 31:533–538.

Awramik, S. M., (1984). Ancient stromatolites and microbial mats. In: Y. Cohen, R. W. Castenholz, and H. O. Halvorson (Eds.), Microbial Mats: Stromatolites. A. R. Liss, N.Y., p. 1-21.

Baumgartner LK, Reid RP, Dupraz C, Decho AW, Buckley DH, Spear JR, et al. (2006). Sulfate reducing bacteria in microbial mats: Changing paradigms, new discoveries. Sediment. Geol. 185:131–145.

Bebout BM, Fitzpatrick MW, Paerl HW. (1993). Identification of the Sources of Energy for Nitrogen Fixation and Physiological Characterization of Nitrogen-Fixing Members of a Marine Microbial Mat Community. Appl. Environ. Microbiol. 59:1495–1503.

Boisvert S, Raymond F, Godzaridis É, Laviolette F, Corbeil J. (2012). Ray Meta: scalable de novo metagenome assembly and profiling. Genome Biol. 13:R122.

Bolger AM, Lohse M, Usadel B. (2014). Trimmomatic: a flexible trimmer for Illumina sequence data. Bioinformatics 30:2114–2120.

Bowman JP, McCammon SA, Lewis T, Skerratt JH, Brown JL, Nichols DS, et al. (1998). Psychroflexus torquis gen. nov., sp. nov., a psychrophilic species from Antarctic sea ice, and reclassification of Flavobacterium gondwanense (Dobson et al. 1993) as Psychroflexus gondwanense gen. nov., comb. nov. Microbiol. Read. Engl. 144:1601–1609.

Brown CT, Sharon I, Thomas BC, Castelle CJ, Morowitz MJ, Banfield JF. (2013). Genome resolved analysis of a premature infant gut microbial community reveals a Varibaculum cambriense genome and a shift towards fermentation-based metabolism during the third week of life. Microbiome 1:30.

Burow LC, Woebken D, Bebout BM, McMurdie PJ, Singer SW, Pett-Ridge J, et al. (2012). Hydrogen production in photosynthetic microbial mats in the Elkhorn Slough estuary, Monterey Bay. ISME J. 6:863–874.

Burow LC, Woebken D, Marshall IPG, Singer SW, Pett-Ridge J, Prufert-Bebout L, et al. (2014). Identification of Desulfobacterales as primary hydrogenotrophs in a complex microbial mat community. Geobiology 12:221–230.

Burow LC, Woebken D, Marshall IP, Lindquist EA, Bebout BM, Prufert-Bebout L, et al. (2013). Anoxic carbon flux in photosynthetic microbial mats as revealed by metatranscriptomics. ISME J. 7:817–829.

Canfield DE, Des Marais DJ. (1991). Aerobic sulfate reduction in microbial mats. Science 251:1471–1473.

Canfield DE, Des Marais DJ. (1993). Biogeochemical cycles of carbon, sulfur, and free oxygen in a microbial mat. Geochim. Cosmochim. Acta 57:3971–3984.

Chang C-C, Lin C-J. (2011). LIBSVM: A Library for Support Vector Machines. ACM Trans Intell Syst Technol 2:27:1–27:27.

Chikhi R, Medvedev P. (2014). Informed and automated k-mer size selection for genome assembly. Bioinformatics 30:31–37.

Cottrell MT, Kirchman DL. (2000). Natural Assemblages of Marine Proteobacteria and Members of the Cytophaga-Flavobacter Cluster Consuming Low- and High-Molecular-Weight Dissolved Organic Matter. Appl. Environ. Microbiol. 66:1692–1697.

Danecek P, Auton A, Abecasis G, Albers CA, Banks E, DePristo MA, et al. (2011). The variant call format and VCFtools. Bioinformatics 27:2156–2158.

Dick G, Andersson A, Baker B, Simmons S, Thomas B, Yelton AP, et al. (2009). Community-wide analysis of microbial genome sequence signatures. Genome Biol. 10:R85.

Dolla A, Fournier M, Dermoun Z. (2006). Oxygen defense in sulfate-reducing bacteria. J. Biotechnol. 126:87–100.

Dupont CL, Rusch DB, Yooseph S, Lombardo M-J, Alexander Richter R, Valas R, et al. (2012). Genomic insights to SAR86, an abundant and uncultivated marine bacterial lineage. ISME J. 6:1186–1199.

Eckburg PB, Bik EM, Bernstein CN, Purdom E, Dethlefsen L, Sargent M, et al. (2005). Diversity of the Human Intestinal Microbial Flora. Science 308:1635–1638.

Eddy SR. (2011). Accelerated Profile HMM Searches. PLoS Comput Biol 7:e1002195.

Ester M, Kriegel H-P, Sander J, Xu X. (1996). A density-based algorithm for discovering clusters in large spatial databases with noise. In: AAAI Press, pp. 226–231.

Everroad RC, Stuart RK, Bebout BM, Detweiler AM, Lee JZ, Woebken D, et al. (2016). Permanent draft genome of strain ESFC-1: ecological genomics of a newly discovered lineage of filamentous diazotrophic cyanobacteria. SIGS. 11:53.

Fike DA, Gammon CL, Ziebis W, Orphan VJ. (2008). Micron-scale mapping of sulfur cycling across the oxycline of a cyanobacterial mat: a paired nanoSIMS and CARD-FISH approach. ISME J. 2:749–759.

Fraser C, Hanage WP, Spratt BG. (2007). Recombination and the Nature of Bacterial Speciation. Science 315:476–480.

Garcia-Pichel F, Prufert-Bebout L, Muyzer G. (1996). Phenotypic and phylogenetic analyses show Microcoleus chthonoplastes to be a cosmopolitan cyanobacterium. Appl. Environ. Microbiol. 62:3284–3291.

Garrison E, Marth G. (2012). Haplotype-based variant detection from short-read sequencing. ArXiv12073907 Q-Bio.

Gill SR, Pop M, DeBoy RT, Eckburg PB, Turnbaugh PJ, Samuel BS, et al. (2006). Metagenomic Analysis of the Human Distal Gut Microbiome. Science 312:1355–1359.

Hanke A, Hamann E, Sharma R, Geelhoed JS, Hargesheimer T, Kraft B, et al. (2014). Recoding of the stop codon UGA to glycine by a BD1-5/SN-2 bacterium and niche partitioning between Alpha- and Gammaproteobacteria in a tidal sediment microbial community naturally selected in a laboratory chemostat. Front. Microbiol. 5.

Harris JK, Caporaso JG, Walker JJ, Spear JR, Gold NJ, Robertson CE, et al. (2013). Phylogenetic stratigraphy in the Guerrero Negro hypersaline microbial mat. ISME J. 7:50–60.

He M, Sebaihia M, Lawley TD, Stabler RA, Dawson LF, Martin MJ, et al. (2010). Evolutionary dynamics of Clostridium difficile over short and long time scales. Proc. Natl. Acad. Sci. 107:7527–7532.

Hess M, Sczyrba A, Egan R, Kim T-W, Chokhawala H, Schroth G, et al. (2011). Metagenomic Discovery of Biomass-Degrading Genes and Genomes from Cow Rumen. Science 331:463–467.

Hyatt D, Chen G-L, LoCascio PF, Land ML, Larimer FW, Hauser LJ. (2010). Prodigal: prokaryotic gene recognition and translation initiation site identification. BMC Bioinformatics 11:119.

Jørgensen BB. (1994). Sulfate reduction and thiosulfate transformations in a cyanobacterial mat during a diel oxygen cycle. FEMS Microbiol. Ecol.13:303–312.

Kashtan N, Roggensack SE, Rodrigue S, Thompson JW, Biller SJ, Coe A, et al. (2014). Single-Cell Genomics Reveals Hundreds of Coexisting Subpopulations in Wild Prochlorococcus. Science 344:416–420.

Khodadad CLM, Foster JS. (2012). Metagenomic and Metabolic Profiling of Nonlithifying and Lithifying Stromatolitic Mats of Highborne Cay, The Bahamas. PLoS ONE 7:e38229.

Langmead B, Salzberg SL. (2012). Fast gapped-read alignment with Bowtie 2. Nat. Methods 9:357–359.

Lee JZ, Burow LC, Woebken D, Everroad RC, Kubo MD, Spormann AM, et al. (2014). Fermentation couples Chloroflexi and sulfate-reducing bacteria to Cyanobacteria in hypersaline microbial mats. Front. Microb. Physiol. Metab. 5:61.

Lee, JZ, D’Haeseleer P, Everroad RC, Prufert-Bebout L, Burow LC, Detweiler AM, Weber PK, Kraoz U, Stuart RK, Lipton MS, Brodie EL, Bebout BM, Pett-Ridge J. Metagenomic analysis of intertidal hypersaline microbial mats from Elkhorn Slough, California, grown with and without additional molybdate. SIGS. (in submission).

Ley RE, Harris JK, Wilcox J, Spear JR, Miller SR, Bebout BM, et al. (2006). Unexpected Diversity and Complexity of the Guerrero Negro Hypersaline Microbial Mat. Appl. Environ. Microbiol. 72:3685–3695.

Li H, Handsaker B, Wysoker A, Fennell T, Ruan J, Homer N, et al. (2009). The Sequence Alignment/Map format and SAMtools. Bioinformatics 25:2078–2079.

Meyer F, Paarmann D, D’Souza M, Olson R, Glass EM, Kubal M, et al. (2008). The metagenomics RAST server – a public resource for the automatic phylogenetic and functional analysis of metagenomes. BMC Bioinformatics 9:386.

Morowitz MJ, Denef VJ, Costello EK, Thomas BC, Poroyko V, Relman DA, et al. (2011). Strain-resolved community genomic analysis of gut microbial colonization in a premature infant. Proc. Natl. Acad. Sci. 108:1128–1133.

Nielsen HB, Almeida M, Juncker AS, Rasmussen S, Li J, Sunagawa S, et al. (2014). Identification and assembly of genomes and genetic elements in complex metagenomic samples without using reference genomes. Nat. Biotechnol. 32:822–828.

Oren A. (2010). Mats of Filamentous and Unicellular Cyanobacteria in Hypersaline Environments. In:Microbial Mats, Seckbach, J & Oren, A, eds (ed). Cellular Origin, Life in Extreme Habitats and Astrobiology, Springer Netherlands, pp. 387–400.

Overmann J, Fischer U, Pfennig N. (1992). A new purple sulfur bacterium from saline littoral sediments, Thiorhodovibrio winogradskyi gen. nov. and sp. nov. Arch. Microbiol. 157:329–335.

Pagani I, Chertkov O, Lapidus A, Lucas S, Del Rio TG, Tice H, et al. (2011). Complete genome sequence of Marivirga tractuosa type strain (H-43T). SIGS. 4:154–162.

Pereira IAC, Ramos AR, Grein F, Marques MC, da Silva SM, Venceslau SS. (2011). A Comparative Genomic Analysis of Energy Metabolism in Sulfate Reducing Bacteria and Archaea. Front. Microbiol. 2.

Podell S, Ugalde JA, Narasingarao P, Banfield JF, Heidelberg KB, Allen EE. (2013). Assembly-Driven Community Genomics of a Hypersaline Microbial Ecosystem. PLoS ONE 8:e61692.

Quinlan AR, Hall IM. (2010). BEDTools: a flexible suite of utilities for comparing genomic features. Bioinforma. Oxf. Engl. 26:841–842.

Rippka R, Deruelles J, Waterbury JB, Herdman M, Stanier RY. (1979). Generic Assignments, Strain Histories and Properties of Pure Cultures of Cyanobacteria. Microbiology 111:1–61.

Rogers TE, Pudlo NA, Koropatkin NM, Bell JSK, Moya Balasch M, Jasker K, et al. (2013). Dynamic responses of Bacteroides thetaiotaomicron during growth on glycan mixtures. Mol. Microbiol. 88:876–890.

Römling U. (2002). Molecular biology of cellulose production in bacteria. Res. Microbiol. 153:205–212.

Ross P, Mayer R, Benziman M. (1991). Cellulose biosynthesis and function in bacteria. Microbiol. Rev. 55:35–58.

Ruvindy R, White Iii RA, Neilan BA, Burns BP. (2016). Unraveling core microbial metabolisms in the hypersaline microbial mats of Shark Bay using high-throughput metagenomics. ISME J. 10:183–196.

Schloissnig S, Arumugam M, Sunagawa S, Mitreva M, Tap J, Zhu A, et al. (2013). Genomic variation landscape of the human gut microbiome. Nature 493:45–50.

Sekiguchi Y, Ohashi A, Parks DH, Yamauchi T, Tyson GW, Hugenholtz P. (2015). First genomic insights into members of a candidate bacterial phylum responsible for wastewater bulking. PeerJ 3:e740.

Sharon I, Morowitz MJ, Thomas BC, Costello EK, Relman DA, Banfield JF. (2013). Time series community genomics analysis reveals rapid shifts in bacterial species, strains, and phage during infant gut colonization. Genome Res. 23:111–120.

Sigalevich P, Meshorer E, Helman Y, Cohen Y. (2000). Transition from Anaerobic to Aerobic Growth Conditions for the Sulfate-Reducing Bacterium Desulfovibrio oxyclinae Results in Flocculation. Appl. Environ. Microbiol. 66:5005–5012.

Spring S, Riedel T. (2013). Mixotrophic growth of bacteriochlorophyll a-containing members of the OM60/NOR5 clade of marine gammaproteobacteria is carbon-starvation independent and correlates with the type of carbon source and oxygen availability. BMC Microbiol. 13:117.

Stuart RK, Mayali X, Lee JZ, Craig Everroad R, Hwang M, Bebout BM, et al. (2015). Cyanobacterial reuse of extracellular organic carbon in microbial mats. ISME J.

Teske A, Ramsing NB, Habicht K, Fukui M, Kuver J, Jørgensen BB, et al. (1998). Sulfate-Reducing Bacteria and Their Activities in Cyanobacterial Mats of Solar Lake (Sinai, Egypt). Appl. Environ. Microbiol. 64:2943–2951.

Varin T, Lovejoy C, Jungblut AD, Vincent WF, Corbeil J. (2012). Metagenomic Analysis of Stress Genes in Microbial Mat Communities from Antarctica and the High Arctic. Appl. Environ. Microbiol. 78:549–559.

Visscher PT, Prins RA, Gemerden H van. (1992). Rates of sulfate reduction and thiosulfate consumption in a marine microbial mat. FEMS Microbiol. Lett. 86:283–293.

Wattam AR, Abraham D, Dalay O, Disz TL, Driscoll T, Gabbard JL, et al. (2013). PATRIC, the bacterial bioinformatics database and analysis resource. Nucleic Acids Res. gkt1099.

Woebken D, Burow LC, Prufert-Bebout L, Bebout BM, Hoehler TM, Pett-Ridge J, et al. (2012). Identification of a novel cyanobacterial group as active diazotrophs in a coastal microbial mat using NanoSIMS analysis. ISME J. 6:1427–1439.

Woyke T, Teeling H, Ivanova NN, Huntemann M, Richter M, Gloeckner FO, et al. (2006). Symbiosis insights through metagenomic analysis of a microbial consortium. Nature 443:950–955.

Wrighton KC, Thomas BC, Sharon I, Miller CS, Castelle CJ, VerBerkmoes NC, et al. (2012). Fermentation, Hydrogen, and Sulfur Metabolism in Multiple Uncultivated Bacterial Phyla. Science 337:1661–1665.

Yurkov V, Stackebrandt E, Holmes A, Fuerst JA, Hugenholtz P, Golecki J, et al. (1994). Phylogenetic Positions of Novel Aerobic, Bacteriochlorophyll a-Containing Bacteria and Description of Roseococcus thiosulfatophilus gen. nov., sp. nov., Erythromicrobium ramosum gen. nov., sp. nov., and Erythrobacter litoralis sp. nov. Int. J. Syst. Bacteriol. 44:427–434.

Zehr JP, Mellon M, Braun S, Litaker W, Steppe T, Paerl HW. (1995). Diversity of heterotrophic nitrogen fixation genes in a marine cyanobacterial mat. Appl. Environ. Microbiol. 61:2527–2532.

Zeng Y, Feng F, Medová H, Dean J, Koblížek M. (2014). Functional type 2 photosynthetic reaction centers found in the rare bacterial phylum Gemmatimonadetes. Proc. Natl. Acad. Sci. 111:7795–7800.

